# *Slit1* -a MET target gene in the embryonic limbs, prevents premature differentiation during mammalian myogenesis

**DOI:** 10.1101/2025.06.22.660860

**Authors:** Masum Saini, Jyoti Jadhav, Giulio Cossu, Sam J. Mathew

## Abstract

Skeletal myogenesis requires precise spatio-temporal regulation of molecular signals during prenatal and postnatal development. Prenatal myogenesis in mouse commences at embryonic day (E) 9.5, characterized by the expression of PAX3 -a key myogenic regulator, and its target MET, in the embryonic muscle progenitor cells (EMPCs). These EMPCs delaminate from dermomyotome in the somites and migrate to designated areas such as the developing embryonic limbs and diaphragm. The trajectory of their migration is directed and limited by spatio-temporal availability of the MET ligand (HGF). Given its periodic expression during embryonic myogenesis and its recent association with familial arthrogryposis, it is important to identify additional non-migratory functions of MET signaling. We find conditional loss of *Met* in the *Pax3*/somitic lineage affects survival in early neonatal stages because amuscular diaphragms cause respiratory distress. Impaired progenitor migration in conditional *Met* knockouts (cMet^KO^) results in highly dysplastic muscles in neonates compared to muscleless limbs observed in previous mutants. Additionally, using cMet^KO^ embryos (E11.5) we identify *Slit1* as a novel target of MET that is downregulated in the mutant limb buds. Pharmacological modulation with SU11274, in vitro, confirms *Slit1* as a MET responsive gene, which if knocked down in myoblasts leads to precocious myogenic differentiation. Similarly, cMet^KO^ embryos, having reduced *Slit1* expression, show greater myotomal differentiation and compact organization, compared to wildtype embryos. While *Slit1* emerges as a MET target that represses precocious switch to myogenic differentiation, molecular intricacies of MET-mediated regulation of *Slit1* and their spatio-temporal dynamic in fine-tuning myogenesis in the nascent limbs needs further examination.

## Introduction

Skeletal muscle is the most abundant mammalian body tissue that can be controlled voluntarily, at variance with cardiac and smooth muscles^1,2^. Mammalian skeletal muscle performs vital functions some of which are posture maintenance, support, motion, metabolism, energy production, homeostasis and pulmonary respiration. Myogenesis, the process of skeletal muscle formation, is intricately regulated in mammals.

Mammalian myogenesis has long been studied using mouse as a model because there is considerable homology between murine and human myogenic genes and transcriptional factors, which allows genetic manipulation of these genes to understand their functional hierarchies and roles in myogenesis^3,4^. In the mouse, myogenesis is accomplished spatio-temporally by sequentially discrete stages –the embryonic, fetal, neonatal and adult myogenesis^2^. While the first two stages occur prenatally, and witness the primary and secondary waves of myogenesis, respectively, the latter two stages comprise postnatal myogenesis^5^. Prenatal myogenesis uses a core set of genetic networks in discrete combinations to establish muscles in distinct body regions^4,6–8^. Many shared elements, such as signaling molecules and transcriptional factors, between pre- and post-natal myogenesis and muscle pathology^4^, suggest the need to further investigate common and distinct mechanisms regulating myogenesis in the embryo and postnatally.

In the mouse embryo, skeletal muscle is established in a multi-step manner where the embryonic phase of primary myogenesis begins first with the formation of the myotomes, patterned stripes of mononucleated, differentiated myocytes around embryonic day (E) 9.5, followed by appearance of small, oligonucleated embryonic or primary myotubes that typically express slow myosin. At around E14.5 fetal myogenesis begins with the formation of multinucleated, larger myotubes that predominantly express fast myosin and mature into cross-striated contractile myofibers. At birth the number of muscle fibers is definitive, and further growth occurs by fusion of still dividing satellite cells^2,4,9^. All skeletal embryonic myogenic progenitor cells (EMPCs) in the body originate from the dorsal somite, the dermomyotome, in a cranio-caudal sequence along the body axis, depending on which, they give rise to specific skeletal muscles in the body^2,4,9^. For example, the EMPCs in the hypaxial dermomyotome of somitic segments adjacent to the limb region undergo lineage commitment, delamination, migrate to and populate the developing limb buds. Physiologically, myogenesis depends upon a fine-tuning between cellular processes of proliferation, differentiation and quiescence ^10,11^.

At the molecular level, myogenesis is orchestrated by a network of signaling pathways with PAX3, a homeodomain transcriptional factor, playing a pivotal role in the early regulatory mechanisms^12^. *Pax3* expression is reported in the pre-somitic paraxial mesoderm even before it is segmented into somites i.e. around E8.5^13,14^. Although induced early in the somites, *Pax3* expression becomes restricted to the central portion of the dermomyotome as the somite matures and persists in the EMPCs that delaminate from the hypaxial dermomyotome to the developing limb buds^8,15,16^. In fact, in Splotch (*Sp*) mice, harboring *Pax3* mutations that result in a functionally null protein, EMPCs fail to migrate to the limb bud^17,18^. Though, PAX3 absence in *Sp* mice is dispensable to differentiation of cells within the myotome of the lateral somite, but in its absence the myotome is disorganized in most mutants^17,18^. PAX3 is also crucial for directly activating *Myf5*, an important myogenic determination factor/muscle regulatory factor (MRF) responsible for lineage commitment in EMPCs of the limb bud^8,19^. Notably, *Sp* mice also lack expression of MET (a proto-oncogenic receptor kinase -RTK encoded by the *c-Met* gene) in the hypaxial dermomyotome, somites and limb buds, which essentially are regions also of PAX3 expression^17,20^. This observation makes it tempting to infer that *Met* is a downstream target of *Pax3*. Though, *Pax3* is known to directly activate *Met*, *in vitro*, it remains to be shown whether *Met* is a direct target of *Pax3 in vivo*^18,21^.

MET is first detected in the ventral region of the somites adjacent to forelimb and hindlimb buds around E9.5 and E10, respectively, which coincides with the first/embryonic wave of myogenesis^18,20^. Upon somitic maturation PAX3+ EMPCs, that are also MET+ now, start delaminating and invading the limb buds in response to hepatocyte growth factor (HGF, which is ligand of MET receptor) that is expressed by the limb bud mesenchyme^20,22–25^. Thus MET signaling causes these cells to undertake long-range migration, resulting in embryonic myoblasts localizing to ventral and dorsal pre-muscle masses in the limbs^20^. The proliferating embryonic myoblasts express MET until E11.5, and eventually fuse with each other to form primary myofibers in the limbs. At E12.5, MET expression is downregulated in primary myofibers in the proximal limb bud region, but its expression is retained distally along the digits. The developing limb muscles at E13.5 show MET re-expression that coincides with renewal of myoblast proliferation, which is required for the ensuing fetal myogenesis. These fetal myoblasts fuse to primary myofibers, which in turn fuse with one another, to form secondary myofibers in the limbs. At E14.5, MET expression in the developing limbs becomes localized in the region of the joints^26^. Recent studies demonstrated that tyrosine residues, which have a crucial role in the activation of MET kinase, harbor mutations that cause muscular dysplasia in patients suffering from familial distal arthrogryposis^27,28^. This condition is characterised by congenital contractures of distal joints due to abnormal development or structure of muscle. These are the first reports of etiological involvement of MET mutations in limb muscle anomaly leading to distal joint contractures in humans.

Recurrent MET expression during pre-natal myogenesis occurs in concurrence with its established roles of promoting progenitor migration and myoblast proliferation^26^. The necessity of MET expression as a signaling molecule that may regulate other genes during myogenesis, remains to be investigated. Therefore, we used conditional *Met* knockout (cMet^KO^) mice, transcriptome sequencing, *in vitro* pharmacological treatment and gene knockdown to explore additional roles of MET in appendicular myogenesis. In agreement with its crucial role in myogenic progenitor migration, we find cMet^KO^ neonates have highly dysplastic appendicular muscles, and amuscular diaphragms that underlie early neonatal mortality. Beyond this important role in EMPC migration, we show that MET signaling regulates *Slit1* expression, which is in turn crucial for normal myogenesis. We find *Slit1* as a novel *Met* target gene that is differentially expressed in cMet^KO^ limb buds. Pharmacological modulation of MET signaling further confirmed that *Slit1* is expressed in a MET-responsive manner. *Slit1* silencing induces premature myoblast differentiation, which is akin to a higher degree of myotomal differentiation and organization seen in cMet^KO^ embryos. In essence, we identify a new function of MET signaling, via its regulation of *Slit1*, of repressing precocious EMPC differentiation as they undertake MET-mediated long range migration during embryonic appendicular myogenesis.

## Results

### Conditional ablation of *Met* results in early neonatal mortality and impacts the formation of appendicular muscle

To investigate additional roles of *Met* in myogenesis, we used a conditional *Met^fl^* allele, where exon 16 (that encodes a critical ATP binding site) in the *Met* gene is flanked by *loxP* sites^29,30^. In E10 embryos, EMPCs migrating to the limb buds are PAX3+ and activate myogenic genes two days after delaminating from the somite. Hence, Pax3 is considered the first molecular marker for this somite-derived migratory EMPC population^14,18,20^. Further, PAX3 and MET show simultaneous spatio-temporal expression in EMPCs of both fore- and hindlimb buds^18,20^.

Therefore, when both alleles *Pax3^CreKI^* and Met^fl/fl^ are combined into one mouse, Cre-recombinase (Cre) expression driven by the endogenous *Pax3* promoter results in a recombined *Met* allele in the EMPCs^31,32^. Consequently, the mutant animal harbors an exon 16 deletion that inactivates the intracellular tyrosine kinase domain, which is crucial to MET-mediated signaling in the targeted lineage/tissue^30,33^. The mating scheme used to generate conditional *Met* knockout (cMet^KO^ = *Pax3^CreKI/+^;Met^fl/fl^*) animals is shown in (Fig. 1A) and deletion of exon 16 was confirmed by PCR genotyping (Fig. 1B). The deletion was also confirmed by real-time quantitative PCR (qPCR) and is described in a subsequent section.

**Figure 1.**
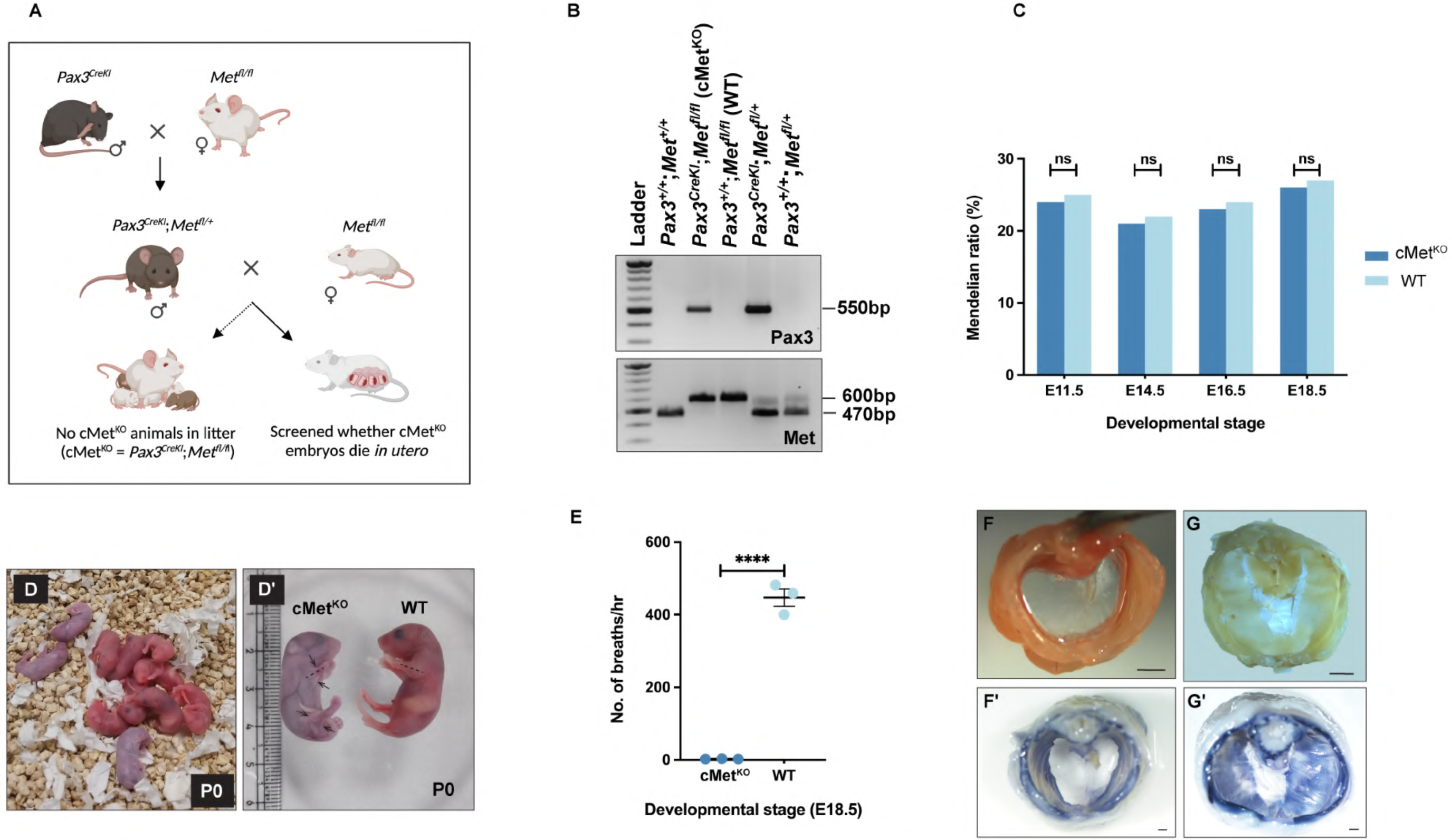
Deleting *Met* in the *Pax3* domain impacts survival. **A.** Breeding scheme to ablate *Met* in the myogenic lineage during development, and generate conditional *Met* knockout (cMet^KO^ i.e. *Pax3CreK^11^+;Met’^1^lfl)* embryos/animals. **B.** PCR-based screening to identify embryos/animals of cMet^KO^ and wildtype (WT) genotypes. **C.** Bar graph showing proportion of cMet^KO^ and *WT* embryos identified at specific developmental stages, where the total number of animals screened is n≥70. Chi-Square test was performed on a contingency table comparing the number of *WT* to cMet^KO^ embryos from timed pregnant females at the indicated developmental stages. **D.** Newborn mouse litter with dead cMet^KO^ and live *WT* pups having cyanotic and bright pink appearance, respectively. **D’.** Comparative muscularization of fore­ and hindlimbs in the cMet^KO^ and *WT* neonates (PO). Arrows indicate loose skin folds in the limbs of the cMet^KO^ neonate. **E.** Breath count data representing the number of breaths per hour (hr) in cMet^KO^ and *WT* animals (E18.5, n=3). The data are represented as means ±SEM and were subjected to unpaired two-tailed T-test. **F-G’.** Brightfield images of PO diaphragm whole mounts, unstained **(F** and G) and stained for myosin (F’ and G’), showing absence of muscle in the cMet^KO^ **(F** and **F’)** compared to *WT* (G and G’) PO pups. Scale bar=1OOOµm.

Intriguingly, no cMet^KO^ animals were found in litters (n=10) that were genotyped around postnatal day (P) 21 (Fig. 1A), suggesting these mice may have developmental defects that do not allow them to survive into adulthood. In mice, developmental defects are known to cause spontaneous embryo resorption during early to mid-gestation (E7-E13) phase and this could possibly explain the absence of cMet^KO^ animals at P21^34,35^. Therefore, using timed matings we screened embryos at specific developmental time-points in both mid (E11.5) and late gestation (E14.5, E16.5 and E18.5). We observed no difference in the Mendelian ratio/proportion of the wildtype (WT = *Pax3^+/+^;Met^fl/fl^*) and cMet^KO^ animals (embryos and fetuses), across the time-points that were screened (Fig. 1C), suggesting that these genotypes survive the perinatal period in utero.

To confirm whether these animals were stillborn or neonatally short-lived, we closely monitored timed pregnant females from E17.5 until birth, to preclude neonatal infanticide and/or cannibalism^36^. We observed the litters had neonates (P0) with both cyanotic and bright pink appearance (Fig.1 D). Genotyping confirmed that the bright pink P0 pups were either WT or non-knockout genotypes, whereas the cyanotic ones were invariably cMet^KO^ mutants. Further, we noted that cyanotic cMet^KO^ pups (P0) rarely showed movement on stimulation and had loose skin folds on the fore- and hindlimbs indicating dysplastic muscles or absence of musculature (Fig.1 D’), which is expected in the cMet^KO^. The cMet^KO^ neonates, if not collected in time were partly or completely cannibalized by the mother, thus explaining their absence at later stages.

### Loss of *Met* in the *Pax3* lineage impinges on survival due to anomalous diaphragm development

Evidently, cMet^KO^ mutants survive in utero, but show cyanosis very early in neonatal life (at P0). Cyanosis in cMet^KO^ neonates indicated oxygen deficiency in the body tissues possibly due to respiratory distress, blood disorders or hypothermia owing to selective neglectful maternal behaviour (may be towards cMet^KO^ pups with defective limb musculature). To confirm the cause of cyanosis we scored the breath counts in cMet^KO^ compared to WT animals.

As a practice, periparturient females are left undisturbed and since the females often give birth at night, it is not always feasible to observe maternal behaviour or document the breathing rates of the newborn pups. Therefore, we harvested E18.5 fetuses from timed pregnant females, which eliminated both -the possibility of maternal behaviour induced respiratory distress and potential loss of neonates to maternal cannibalism. cMet^KO^ mutants were visually distinguishable from the non-cMet^KO^ fetuses, based on morphology of their fore- and hindlimbs. Since the cMet^KO^ mutants were neonatally short-lived, the genotypes were re-confirmed by genotyping tail snips after recording the breath counts. There was a considerable difference in the breathing rates (breath count/hour) between the cMet^KO^ and non-knockout animals (Fig. 1E), which is documented also in a representative two minute clip (Supplementary Video 1) of recordings used for Fig. 1E. Clearly, breathing in cMet^KO^ is compromised resulting in respiratory distress, which explains the cyanosis in cMet^KO^ neonates.

To identify the underlying cause of respiratory distress in cMet^KO^ neonates, we first tested the possibility of a developmental defect in the diaphragm. This is because, PAX3 expressing EMPCs colonise the developing diaphragm around E10.5, and *Met* is known to have crucial role also in diaphragmatic muscularization. Therefore, an absence of either *Pax3* or *Met* leads to impaired diaphragmatic development^22,37,38^. We indeed observed a lack of muscularization, validated by the absence of myosin staining in the diaphragm of cMet^KO^ mutants (Fig. 1F and 1F’) compared to the WT P0 neonates (Fig. 1G and 1G’). This diaphragmatic defect underlies the distressed breathing and compromised oxygenation leading to cyanosis and ensuing mortality in the cMet^KO^ neonates at P0.

### Conditional *Met* mutants show aberrant myogenesis in limb skeletal muscles

The cMet^KO^ pups (P0) showed substantially reduced body weight compared to WT neonates, which is consistent with the altered phenotype of limb muscles seen in the cMet^KO^ mutants (Fig. 2A and Fig. 1D’). Deskinning and staining the fore- and hindlimbs of P0 pups, for myosin, showed diffuse, possibly non-specific residual staining in cMet^KO^ P0 mutants (Fig. 2B and 2C) opposed to a normally patterned myosin staining in the WT neonates (Fig. 2B’ and 2C’). Neurofilament staining was seen in fore- and hindlimbs of both the cMet^KO^ (Fig. 2D and 2E) and WT neonates (Fig. 2D’ and 2EC’), which is in line with previous observation^38^. However, it needs to be ascertained whether absence of skeletal muscle alters pattern or functional aspects of nerve innervation of the limbs. Non-specific neurofilament staining of blood vessels in the hindlimb was ruled out by performing a no primary antibody control (Supplementary Fig. 1B and 1B’).

**Figure 2.**
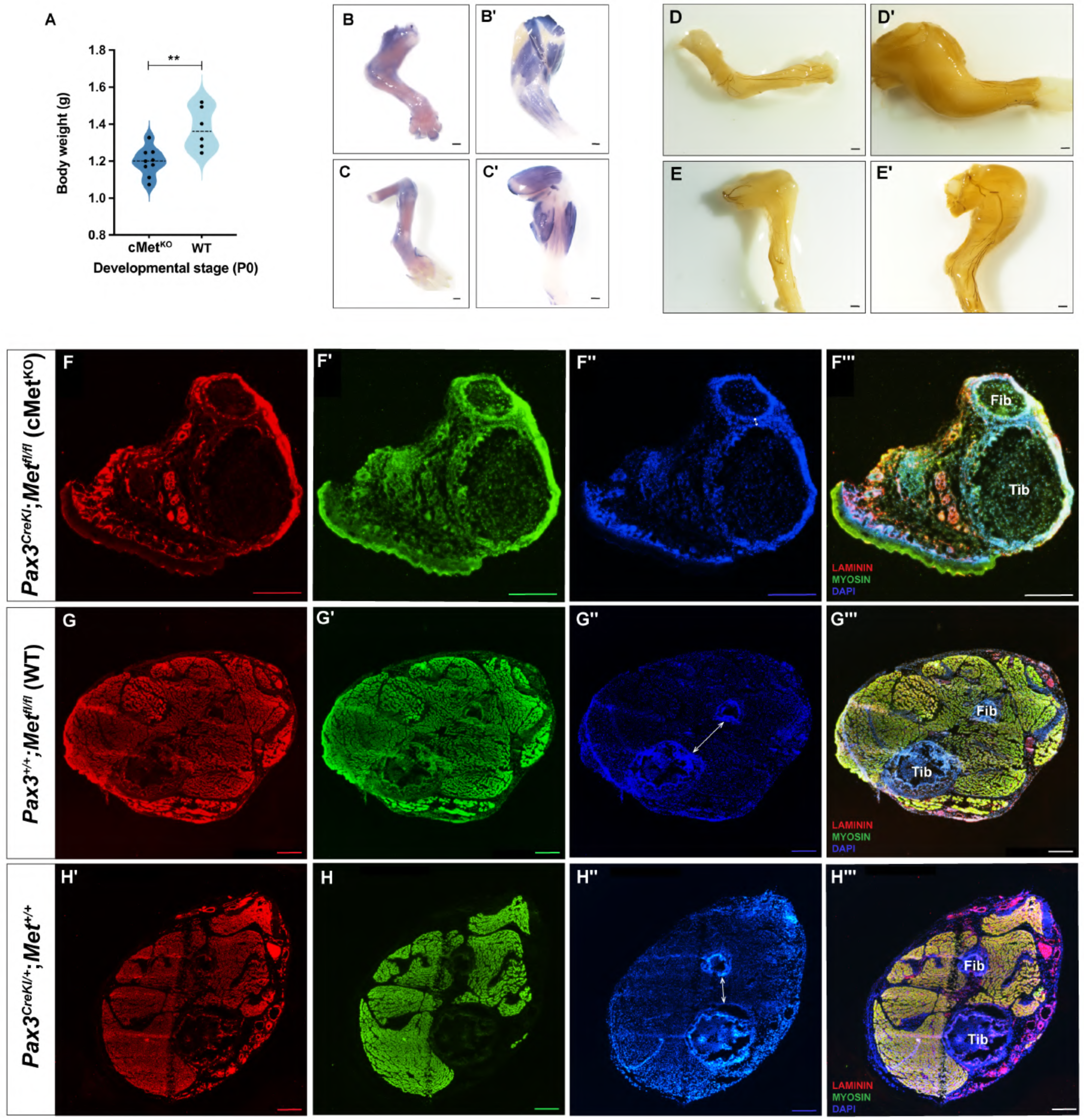
Conditional *Met* knockout animals show impaired skeletal myogenesis in limbs. A. Bar graph depicting body weight of conditional *Met* knockout (cMet^KO^) and wildtype (WT) neonates (PO, n≥6). The data was analysed using unpaired two-tailed T-test. **_B-C’._** Whole-mount myosin staining (with AP) of fore- and hindlimbs from cMet^KO^ (B and C) and WT (B’ and C’) PO neonates, respectively. Scale bar=10OOµm, AP=alkaline phosphatase. **D-E’.** Whole-mount neurofilament staining (DAB) of fore- and hindlimbs from cMet^KO^ (D and D’) and WT (E and E’) neonates (PO), respectively. Scale bar=1OOOµm, DAB=diaminobenzidine. F-H”’. Transverse sections through hindlimbs of cMet^KO^ (F-F”’), WT (G-G”’) and *_Pax3creKI_* (H-H”’) neonates (PO), stained for myosin (green) and laminin (red), and nuclei stained with DAPI (blue). Double-headed white arrows in F”, G”, and H”, indicate tibio-fibular distance. Scale bar=2OO µm, Tib=Tibia, Fib=Fibula.

To rule out residual myosin staining seen in cMet^KO^ fore- and hindlimbs as an artefact, we cross-sectioned the hindlimbs of cMet^KO^ and WT neonates (P0) and stained with myosin (green) and laminin (red) antibodies, and DAPI (blue). We observed random myosin staining resembling disorganized abortive myofibers loosely arranged within the cMet^KO^ hindlimb sections (Fig 2F-F’’’), suggesting dysplastic myogenesis opposed to completely amyogenic limbs in the complete/constitutive *Met* mutants. The myosin staining showed normally organized myofibers in the cross-sections of hindlimbs from both WT (Fig 2G-G’’’) and *Pax3^CreKI^* (Fig 2H-H’’’) neonates (P0). Also, cMet^KO^ hindlimb cross-sections (Fig 2F’’) showed reduced separation between tibia and fibula compared to WT and *Pax3^CreKI^*(Fig 2G’’ and 2H’’). This observation is in accordance with previously documented reduction in tibio-fibular distance in Myf5^-/-^ and Myf5^-/-^:Myod^-/-^ mutant hindlimbs that lack limb musculature^39^.

### Genes of the skeletal muscle biology pathways are differentially expressed in *Met* knockout embryonic limb buds

Migration of EMPCs in the forelimb buds is considered complete by embryonic stage E10.5 but is still underway in the hindlimb buds^40,41^. *Met* continues to be expressed in proliferating embryonic myoblasts at E11.5^26^. Thus, E11.5 seemed an appropriate stage to study the non-migratory roles of MET signaling during appendicular myogenesis. Accordingly, we used fore- and hindlimb buds from E11.5 embryos of the cMet^KO^ and WT genotypes, for RNA sequencing (Fig. 3A). The Venn diagram shows differentially expressed genes (DEGs) between the compared genotypes, that overlap or are exclusive to the fore- and hindlimbs (Fig. 3A).

**Figure 3.**
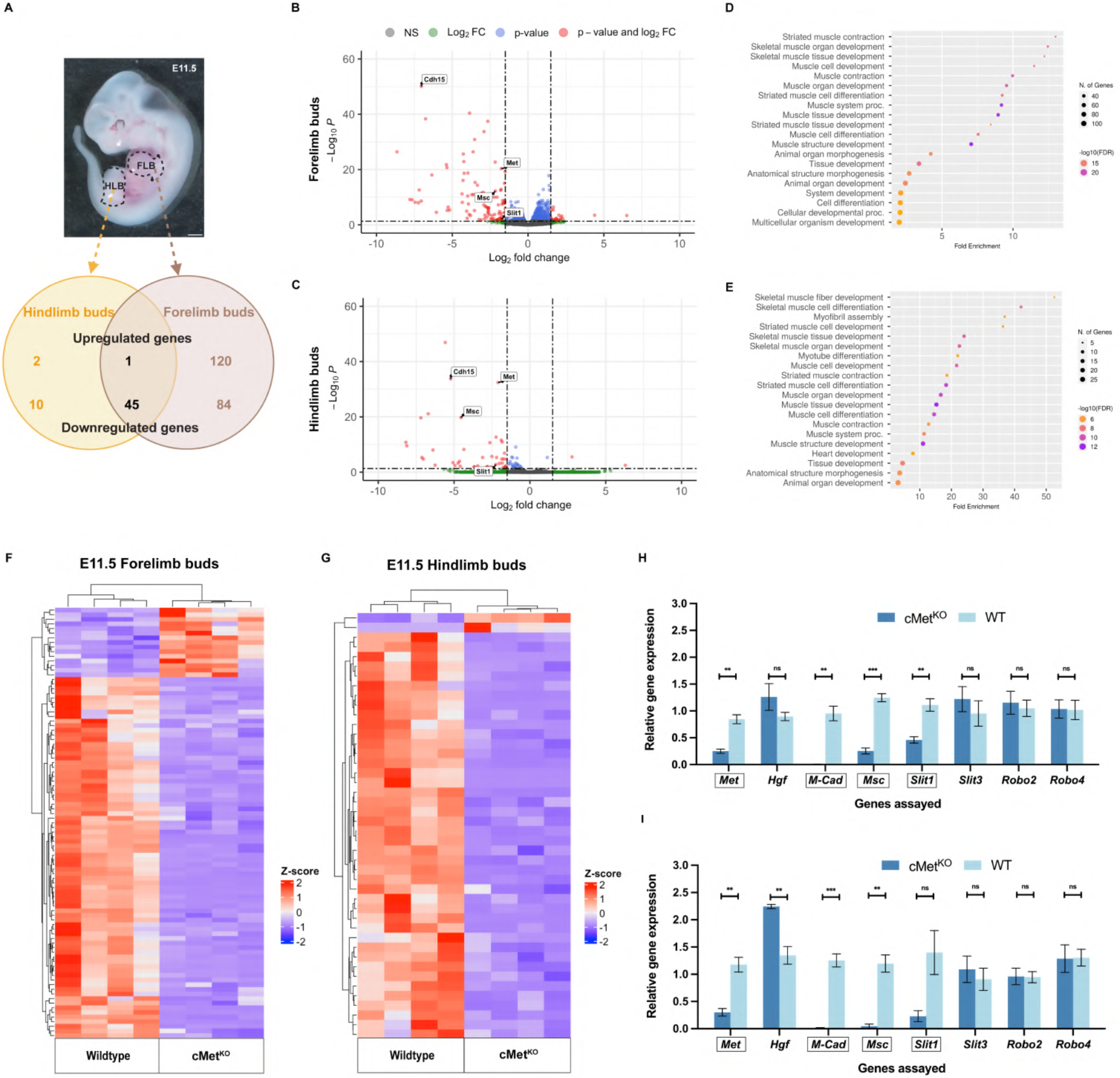
Transcriptome analyses of limb buds between wildtype and conditional *Met* knockout embryos at E11.5, identifies *Slit1* as a target. **A.** A representative whole-mount brightfield image of an E11.5 mouse embryo with FL8 and HL8 marked with a dotted black line. Scale bar=1000µm. A pair of FL8s and HLBs from four embryos of each genotype (cMet^KO^ and WT) were harvested, and subjected to RNA-sequencing analysis. Venn diagram depicts both significantly upregulated and downregulated DEGs that overlap between FL8s and HL8s, in E11.5 mouse embryos. **B and C.** Volcano plots representing gene expression changes in FL8s (A) and HL8s (8) of cMet^KO^ versus WT embryos (E11.5). **D and E.** Dot plots representing Gene Ontology (GO) pathway enrichment analysis of DEGs identified by RNAseq. The top 20 most significant pathways in the GO biological process enrichment results, in FL8s (A) and HL8s (8) from the two compared groups (cMet^KO^ and WT E11.5 embryos) are shown. F **and G.** Heatmaps showing 2-dimensional hierarchical clustering with Z-score scaling method to cluster genes with similar expression profiles between samples. **H and** I. _Quantifying expression of representative genes in both FL8s (H) and HLBs (I) between the compared groups (cMet^KO^ and WT E11.5 embryos) by quantitative real-time PCR (qPCR). The qPCR data validated the genes (boxed) that were downregulated in RNAseq. qPCR was performed using an aliquot of the same RNA from FL8 and HL8 replicates that were previously used for RNA sequencing. The qPCR data is represented as means ±SEM and were subjected to multiple unpaired T-tests. Cdh15 gene (B and C) is also known as M-Cadherin (M-Cad) (H and I). FL8s *=* Forelimb buds, HL8s *=* Hind limb buds, ns=non-significant._

RNAseq data from the forelimb buds (FLBs) and hindlimb buds (HLBs) were assessed for its quality (Supplementary Fig. 2) and is represented by volcano plots that depict the effect of conditional *Met* ablation on gene expression by stratifying gene hits into specific categories (Fig. 3B and 3C). In the volcano plots the X-axis (Log_2_ fold change) and Y-axis (-Log10 of P value) indicate fold change (FC) in gene expression and the significance level of the difference in gene expression, respectively, in the compared groups (cMet^KO^ and WT). Red dots depict genes that are significantly altered (Log_2_ fold change ≥ 1, or ≤ −1, and P<0.05). Red dots towards the positive and negative side of the X-axis indicate significantly upregulated and downregulated genes (i.e. DEGs), respectively. Blue dots show genes that have more than or equal to two-fold change in expression, but P≥0.05. Green dots represent genes where P<0.05 but change in expression (Log_2_ fold change) is less than two-fold. Grey dots depict genes that have less than two-fold change and P≥0.05 and are thus statistically non-significant (NS). Based on these filtering criteria, HLBs show lesser number of both downregulated and upregulated DEGs in contrast to the FLBs. Some of the DEGs indicated by connectors and gene labels on the volcano plots, were validated further as they were downregulated in both the FLBs (Fig. 3B) and HLBs (Fig. 3C) between the compared genotypes and were also enriched in the gene ontology enrichment analysis described below.

The DEGs uncovered by RNAseq were subjected to a gene ontology (GO) enrichment analysis to identify significantly enriched biological processes. In the GO enrichment plots (Fig. 3D and 3E), X-axis ‘Fold Enrichment’ represents magnitude of enrichment i.e. ratio of DEGs enriched in a pathway to the total genes in that pathway, and higher the value stronger the enrichment of the pathway. The Y-axis indicates significant pathways that are filtered first by the false discovery rate (FDR) value and then ranked by fold enrichment. The FDR is a ratio of the false positive results to the total positive test results. The lower the FDR-adjusted P-value, also called FDR q-value (typically below 0.05) indicates a lower likelihood that enrichment is a chance event. - Log_10_(FDR) is the -Log_10_ transformation of FDR q-values. A lower -Log_10_(FDR) value indicates higher significance of enrichment.

A dot or bubble in the plot represents the genes enriched in the corresponding pathway. The distance of the dot along the X-axis indicates the magnitude of enrichment of the corresponding biological pathway. The size of the dot indicates the number of genes in the pathway that overlap with the probed DEGs. The colour of the dot corresponds to a -log10(FDR) value and represents the enrichment confidence or statistical significance. Typically, a lower value indicates more significant enrichment. The pathways related to skeletal muscle development, differentiation, structure and contraction, and to animal organ development and morphogenesis were significantly enriched in RNAseq datasets of FLBs (Fig. 3D) and HLBs (Fig. 3E) (E11.5 embryos). A comparative account of differential gene expression in FLBs (Fig. 3F) and HLBs (Fig. 3G) of cMet^KO^ E11.5 embryos compared to those of the WT genotype, has been rendered using heat maps. The samples (X-axis) and DEGs (Y-axis) are clustered based on the similarity of their Z-scores. The Z-score (colour scale) of each gene quantifies the number of standard deviations a data point (gene expression value) is away from the mean of a dataset (corresponding row or column). Based on the enrichment analysis we picked M-Cadherin (*Cdh15*/*M-Cad*), Musculin (*Msc*) and *Slit1* as candidates, which represented also in the volcano plots described above (Fig. 3C-B).

To validate and analyse the 3 candidate genes we first quantified their expression, along with expression of some other genes by qPCR, using aliquots of the same RNA samples from the 4 replicate FLBs and HLBs, which were used for RNA sequencing. In agreement with its gene ablation, *Met* expression was significantly downregulated in the FLBs (Fig. 3H) and HLBs (Fig. 3I) of cMet^KO^ E11.5 embryos compared to the WT ones. The *Slit1* transcript levels show a striking downregulation of nearly 6-fold in the cMet^KO^ compared WT HLBs (Fig. 3I). Nevertheless, this difference may seem statistically non-significant owing to a large variation in the *Slit1* levels in WT HLB replicates. Overall, the qPCR-based quantitative analysis validated reduced expression of genes, such as *M-Cad*, *Msc* and *Slit1* in FLBs (Fig. 3H) and HLBs (Fig. 3I), that were downregulated also in the RNAseq data (Fig. 3B, 3C, 3F and 3G).

### Pharmacological modulation of MET signaling *in vitro*, identifies *Slit1* as a downstream target in myoblasts

Since, expression of *M-Cad*, *Msc* and *Slit1* transcripts was significantly downregulated in FLBs and HLBs from cMET^KO^ embryos these genes could be downstream targets of MET signaling. M-CAD is a skeletal muscle-specific transmembrane protein that mediates cell-cell adhesion during prenatal and postnatal myogenesis and is a transcriptional target of PAX3 and MyoD^42–47^.

MSC is a basic helix-loop-helix (bHLH) transcriptional factor and a repressor of myogenesis during embryonic stages and in muscle regeneration^48–50^. Its expression in the non-myogenic lineages and muscle precursors is inversely associated with the extent of differentiation, which is considered vital in repressing myogenic differentiation program and preventing premature or improper muscle formation^51^. *Slit1* belongs to a family of 3 homologs (*Slit1-3*), which encode secreted proteins/ligands that bind to specific cognate Roundabout (Robo) receptors encoded by family of 4 homologous genes (*Robo1-4*). Slit-Robo signaling plays crucial role in -axon guidance during development of nervous system, and in the development lung, kidney and mammary gland^52,53^. SLIT1 is crucial for migration of muscle progenitors/myoblast pioneers expressing Robo2 during myotomal patterning in the chick embryo, but its role in murine limb skeletal myogenesis is not known^54^.

Based on the involvement of the 3 candidate genes in the myogenic program and their downregulation observed in limbs of cMet^KO^ embryos (having conditionally inactivated MET kinase), we tested whether MET signaling regulates these genes in the murine C2C12 myoblasts using pharmacological activation and inhibition. Binding of HGF (MET ligand), to MET activates its kinase activity which in turn transduces downstream signaling through different pathways including the mitogen-activated protein kinase (MAPK) cascade, phosphoinositide 3-kinase–Akt (PI3K–Akt)^55^. These pathways mediate MET-dependent cell migration, proliferation, survival and differentiation^55^. They are known to be inhibited by SU11274, a known inhibitor of MET phosphorylation^56^. Accordingly, we assayed expression of the three genes in proliferating C2C12 cells in response to a) stimulation of MET signaling by HGF, b) its inhibition by SU11274, and c) sequential treatment with SU11274 and then HGF.

Representative immunoblots showing post-treatment protein levels and bar graphs illustrating corresponding densitometric analysis of the three replicates, are presented in Fig. 4 Expectedly, levels of phosphorylated (P)-MET (at tyrosine residues 1234/1235) (Fig. 4A and 4A’), SLIT1 (Fig. 4B and 4B’) and P-p44/42 (Fig. 4B and 4B’’), M-CAD (Fig. 4C and 4C’) and P-Akt (Fig. 4C and 4C’’), and MSC and P-p44/42 (Supplementary Fig. 1), show a discernible decrease in response to inhibition by SU1127 compared to HGF-mediated activation. Next, we used specific inhibitors to identify downstream pathways that exert MET signaling-mediated regulation. U0126 inhibits activation of extracellular signal-regulated kinase (ERK1/2) in the MAPK pathway, while Wortmannin (WO) inhibits Akt phosphorylation^57,58^. We followed the same pharmacological treatment strategy, as in the case of SU11274-mediated inhibition, and examined candidate gene expression in proliferating C2C12 cells. In coherence with their mechanism of action, U0126 and WO inhibited phosphorylation of p44/42 (i.e. ERK1/2, (Fig. 4D and 4D’’) and Akt (Fig. 4E and 4E’’), respectively. Further, we observed that SLIT1 levels were downregulated by inhibition of MAPK pathway (Fig. 4D and 4D’), whereas M-CAD expression was modulated by AKT signaling (Fig. 4E and 4E’).

**Figure 4.**
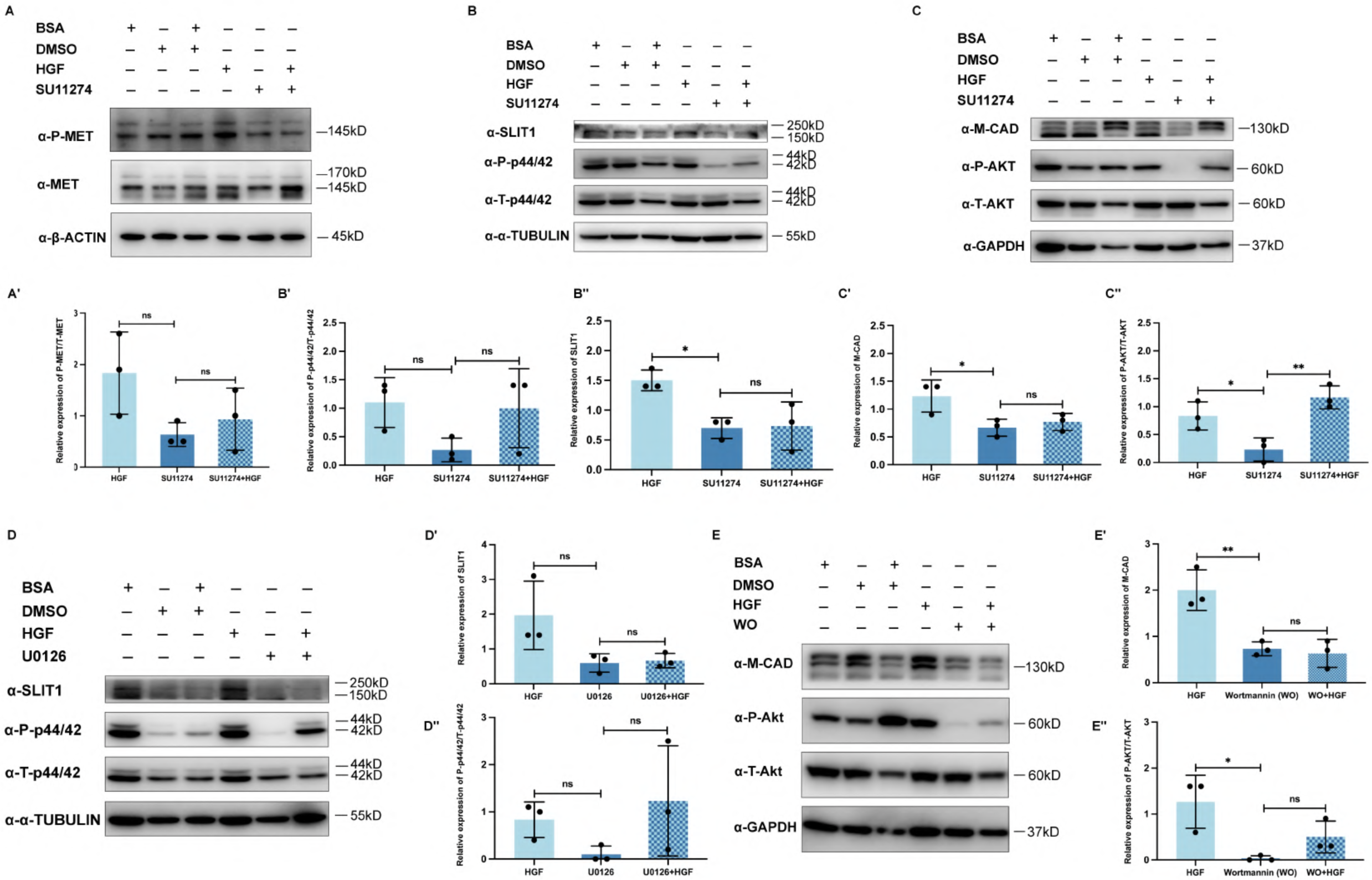
MET signaling modulates the expression of candidate genes in mouse myoblasts. A-C.Representative immunoblots using lysates of C2C12 cells, showing levels of phosphorylated (P)-MET tyrosine 1234/1235 and total (T)-MET (A), SLIT1, P-p44/42 and T-p44/42 (8), and M-CAD, P-Akt and T-Akt (C), in response to HGF-mediated stimulation, inhibition by SU11274, sequential treatment with SU11274 and HGF, and treatment with respective solvents/vehicles in which the pharmacological reagents were dissolved. **A’-C”.** Bar graphs showing densitometric quantitation of P-MET 1234/1235 (A’), SLIT1 (B’), P-p44/42 (B”), M-CAD (C’) and P-Akt (C”) using the expression data from immunoblots (A-C). **D-D”.** Representative western blots and corresponding bar graphs showing densitometric values of SLIT1 (D and D’) and P-p44/42 (D and D”) levels upon treatment of C2C12 cells with HGF, U0126, U0126+HGF, and with respective solvents of these reagents. **E-E”.** Representative immunoblots and corresponding densitometry data graphs showing levels of M-CAD (E and E’) and P-Akt (E and E”) in C2C12 cells treated with HGF, Wortmannin (WO), WO+HGF and their respective solvents. Densitometry data in the bar graphs represent treatment-responsive protein expression normalized initially to the reference gene and then to protein levels in the respective vehicle/solvent controls. Graphical data are represented as means ±SD and significance was tested using one-way ANOVA. _37_

We observed, HGF treatment post-U0126 and WO-mediated inhibition rescued expression of P-p44/42 (Fig. 4D’’) and P-Akt (Fig. 4E’’), respectively. Similar trend of restoration in levels of P-MET (Fig. 4A’), P-p44/42 (Fig. 4B’’) and P-Akt (Fig. 4C’’), in response to HGF treatment post-SU11274 inhibition, was observed. However, in certain instances (Fig. 4A’, 4B’, 4B’’, 4D’’ and 4E’’) the rescue seemed statistically non-significant, which was due to notable variation between replicates. Similarly, inhibitor-mediated decrease in expression of P-MET (Fig. 4A’), SLIT1 (Fig. 4D’) and P-p44/42 (Fig. 4B’’ and 4D’’) though notable remained statistically irrelevant because replicates in their corresponding HGF-treated groups were not tight. When replicates were tightly grouped, such as in case of P-Akt levels (Fig. 4C’’), inhibition and rescue were significant. Notably, there was a marginal rescue in SLIT1 expression upon HGF treatment post-inhibition with SU11274 (Fig. 4B’) and U0126 (Fig. 4D’). Similarly, M-CAD expression was minimally rescued by HGF treatment post-SU11274 (Fig. 4C’) and WO (Fig. 4E’) inhibition. Although, we have not examined this further, it may be due to several factors: a) inability of HGF to override the effect of inhibitors at the concentration used or duration of treatment, b) translational lag resulting in delayed accumulation of proteins to reflect a considerable rescue, c) possible auto regulatory feedback loop in case of SLIT1 ligand which may take time to kick-in and restore its levels, d) inhibition of other downstream pathways transduced by MET such as p38 MAPK, which may regulate *Slit1* and *M-Cad* but become blocked because of SU11274 treatment, and e) in case of U0126 and WO, an upstream activation of MET by HGF treatment is counteracted by these inhibitors as they act on pathways downstream to MET and hence produce an attenuated rescue of candidates. Overall, the considerable decrease in candidate gene expression upon inhibition by SU11274, and in response to pathway-specific inhibitors suggests these genes are regulated in a MET signaling-responsive manner, which is mediated by the ERK-MAPK and Akt pathways in case of SLIT1 and M-CAD, respectively.

### Knocking down *Slit1* expression induces premature myogenic differentiation in myoblasts

In the developing chick embryo, Slit1 is a sclerotomal signal that interacts with its receptor Robo2 on the myoblasts, and this interaction is crucial for myotomal morphogenesis^54^. It has been reported that MET signaling is active in the murine somitic myogenic cells, but not in the limb fibroblasts^38^. We identified *Slit1* as a DEG in an RNA sequencing analysis of the developing mouse limb buds following the somitic ablation of *Met*. Additionally, we observed that its expression varies in a MET signaling-responsive manner in C2C12 cells. These findings imply that in mouse, SLIT1 is produced by the myogenic cells. To test its function in myogenesis we knocked down *Slit1* expression in the proliferating murine C2C12 myoblasts that were cultured in pro-myogenic conditions. Remarkably, compared to control siRNA transfected C2C12 cells which typically form myofibers around day five of differentiation (D5) (Fig. 5A-A’’’), downregulation of *Slit1* expression led to premature fusion and differentiation of myoblasts into myofibers by D2 (Fig. 5B’ and Supplementary Fig. 1C),

**Figure 5.**
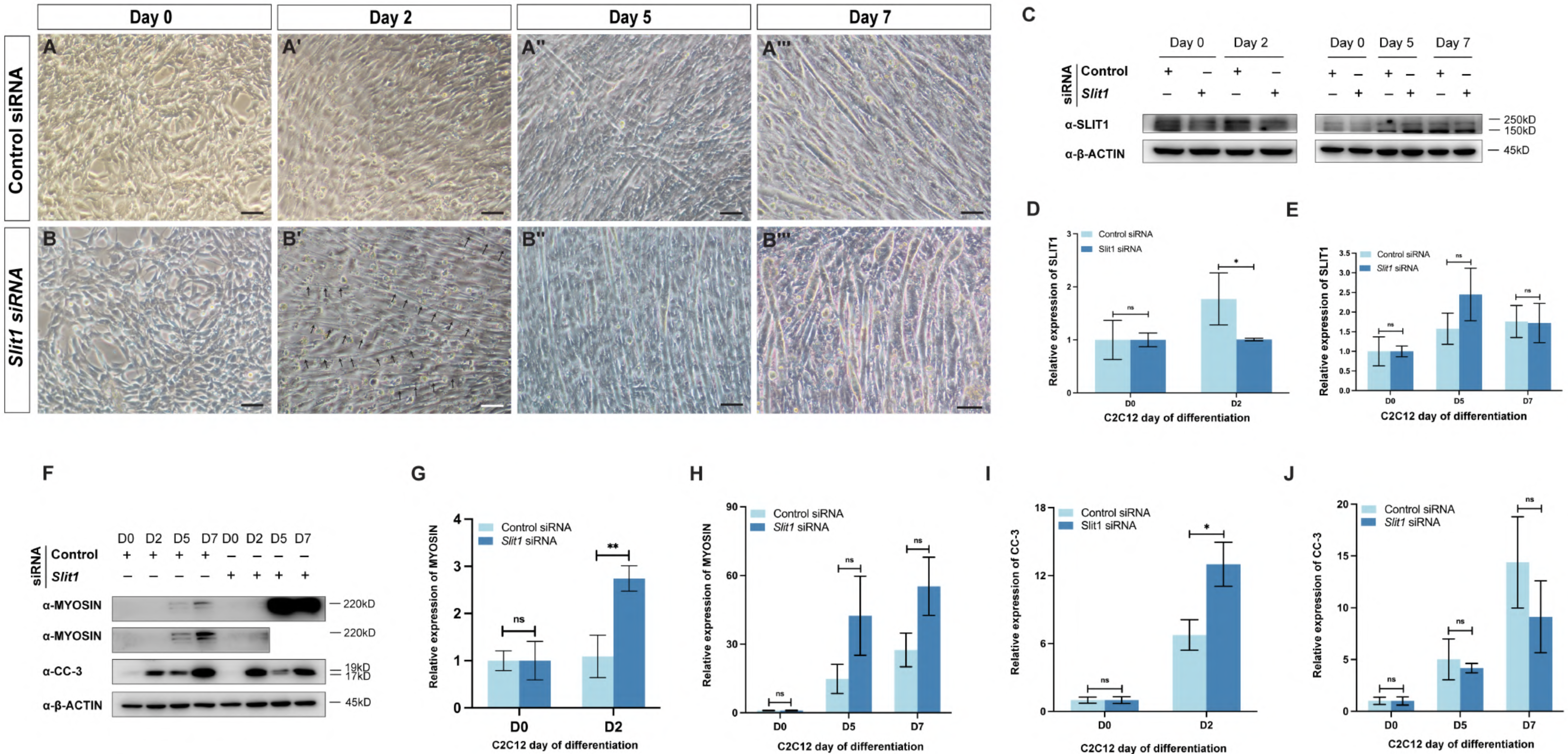
Silencing *Slit1* in murine myoblasts leads to pre-mature differentiation. **A-A”’.** Control siRNA transfected C2C12 mouse myoblasts show myotube formation by Day (D) 5 of differentiation protocol. **B-B”’.** siRNA-mediated downregulation of *Slit1* in C2C12 cells leads to the formation of myofiber like structures, highlighted by black arrows (B’), quite premtaurely i.e. on second day (D2) of differentiation. **C.** Representative immunoblots showing levels of SLIT1 and ≥-ACTIN, at DO, D2, D5, and D7 of myoblast differentiation, in cell lysates of C2C12 cells transfected with control or *$/it1* siRNA. D-E. Bar graphs representing densitometric analyses of SLIT1 expression at DO and D2 (D), and at DO, D5, and D7 (E) of differentiation, in control versus *Slit1* siRNA transfected C2C12 lysates as represented in the corresponding immunoblots (C). **F-1.** Representative western blots (F) and their densitometric quantitation (G-J), depicting expression of Myosin (F-H) and Cleaved Caspase 3 (CC-3) (F, 1-J) at indicated day (D) of myogenic differentiation in lysates from C2C12 cells transfected with control or *Slit1* siRNA. For densitometric quantification, SLIT1, Myosin and CC-3 expression was normalized to corresponding ≥-ACTIN levels at each differentiation day for both (control and *$/it1)* siRNA transfections. Expression of gene of interest, normalized to reference gene values at indicated days of differentiation, were in turn normalised to similarly normalized DO values of the respective siRNA transfection. The graphical data are represented as means +SD and were subjected to multiple unpaired T-tests. ns=non-significant.

We validated downregulation of *Slit1* expression in C2C12 cells by immunoblotting (Fig. 5C). Notably, SLIT1 expression was considerably reduced at D2 in *Slit1* versus control siRNA transfected cells (Fig. 5D). However, this repression was lost around D5 and onwards (Fig. 5E). Nonetheless, since differentiation was already set in motion, the myoblasts transfected with *Slit1* siRNA maintained their myogenic edge compared to the control cells. This is evident from the remarkably high levels of Myosin expression at D2 (Fig. 5F and 5G), which remained discernibly higher through the later days of differentiation (Fig. 5F and 5H) in *Slit1* siRNA transfected cells versus the control cells. The first and second immunoblot images showing Myosin levels (Fig. 5F) were imaged using the same membrane/blot. The blot in the second image was subjected to a longer exposure duration and by concealing last two lanes corresponding to D5 & D7 of *Slit1* siRNA cell lysates, which had in the first image prevented the development of bands in lanes with lower Myosin expression i.e. D2 of *Slit1*, and D5 and D7 of control siRNA transfected cells. To overcome this issue, we electrophoresed and probed D0 and D2 lysates from control and *Slit1* siRNA transfected cells, on a separate gel and membrane, respectively. A distinct Myosin band was detected at D2 of differentiation only in the *Slit1* siRNA transfected cells (Supplementary Fig. 1D). Notably, cleaved caspase 3 (CC-3) levels, a marker of cell death, were elevated at D2 in *Slit1* versus control siRNA transfected C2C12 cells (Fig. 5F and 5I), which is consistent with increased cell death at onset of myogenic differentiation and during subsequent fusion of cells to form myotubes^59^. Accordingly, CC-3 levels increased in control cells as they started to fuse, forming myofibers and myotubes D5 onwards (Fig. 5F and 5J). Increase in CC-3 levels at D7 compared to D5 in *Slit1* siRNA transfected samples (Fig. 5F and 5J) is in agreement with myotube detachment seen in later stages of differentiation^59^. Thus, *Slit1* silencing causes precocious myogenic differentiation of murine myoblasts which is accompanied by molecular hallmarks of myogenesis.

### Somitic myotome is more differentiated in conditional *Met* knockout embryos, phenocopying *in vitro* silencing of *Slit1* in myoblasts

The data so far, demonstrate that MET signaling regulates *Slit1* expression, which is in turn crucial for preventing premature myogenesis. Therefore, it is reasonable to hypothesize that MET signaling inhibits premature differentiation of EMPCs as they migrate into the developing limb buds. To test this we examined the mutant embryos where EMPCs, because of their recombined *Met* allele, are signaling deficient and have altered levels of Slit1 expression.

First, we immunostained transverse sections of cMet^KO^ and WT embryos (E11.5) through the limb bud region using antibodies against MET (red) and PAX3 (green), and stained the nuclei with DAPI (blue). The anti-MET antibody used for immunostaining recognises an epitope in the extracellular region of the RTK, which is outside the region knocked out by Cre-mediated recombination, and hence detects the mutant protein. In agreement with the known role of MET signaling in EMPC migration, cMet^KO^ embryos did not show pre-muscle masses on the dorsal and ventral sides of FLBs (Fig. 6A-6A’’’) and HLBs (6C-6C’’’) compared to WT embryos (Fig. 6B-6B’’’ and 6D-6D’’’) at E11.5. These pre-muscle masses, formed by migrated EMPCs, largely comprised MET+ cells, interspersed with MET and PAX3 co-stained cells (MET+;PAX3+ cells). The MET+;PAX3+ cells were also present outside the pre-muscle masses, along the migratory path in the limb buds. In addition to the absence of pre-muscle masses, cMet^KO^ E11.5 embryos showed considerably reduced numbers of MET+;PAX3+ EMPCs in FLBs (Fig. 6E) and HLBs (Fig. 6F), compared to WT embryos. It is known that embryonic progenitors in mouse embryos (E10.5) lacking *Met* or *Hgf* are properly specified, but fail to migrate and remain aggregated close to the somite^23^. However, at E11.5 we did not notice any conspicuous aggregates of either MET or PAX3 immunopositive cells close to the somite in the cMet^KO^ embryos, suggesting that either these cells apoptose or commit to the myogenic lineage fusing with the myocytes of the somitic myotome that is undergoing myogenic differentiation.

**Figure 6.**
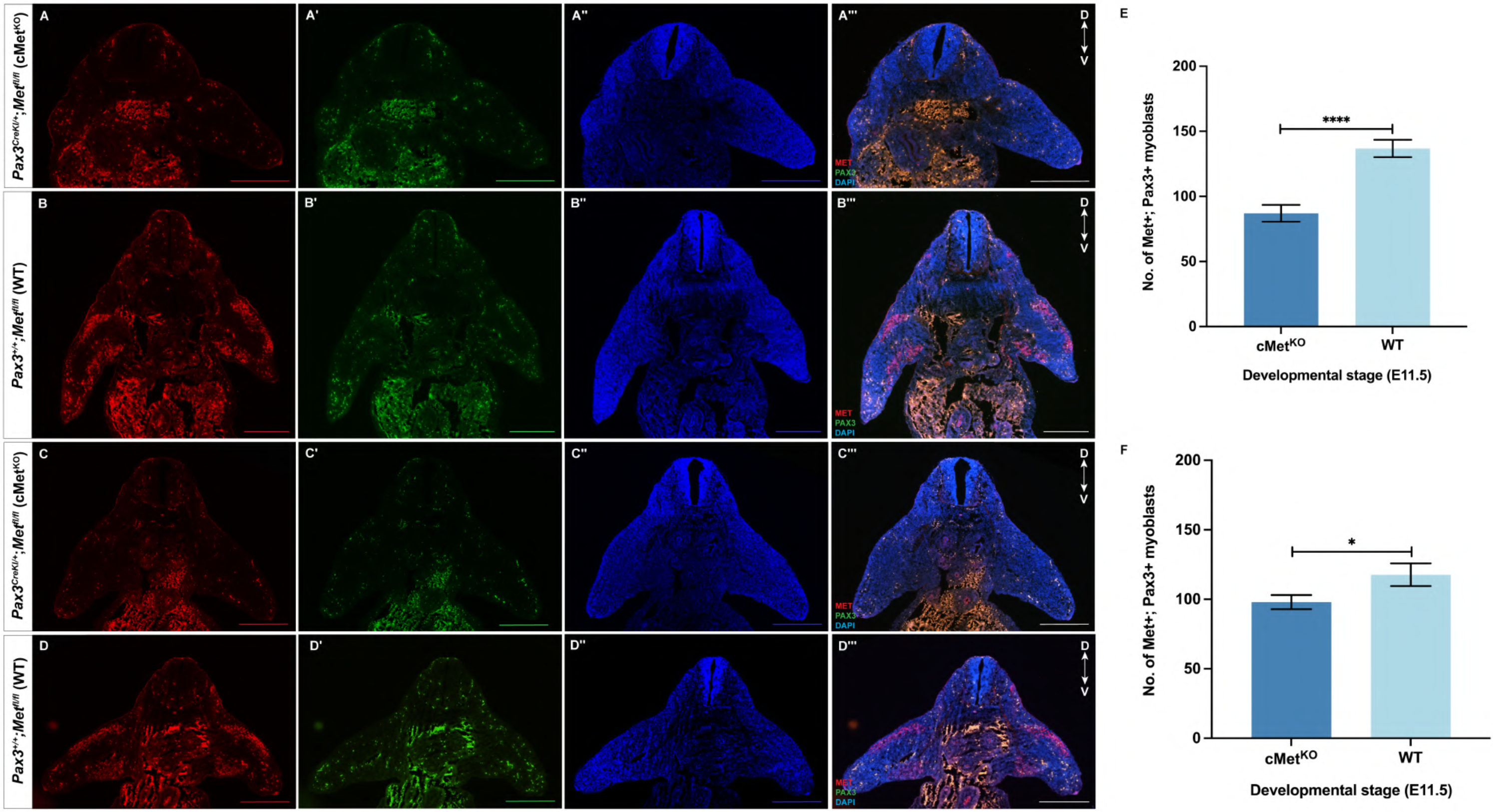
Developing limbs in conditional *Met* knockout embryos lack pre-muscle masses and have a reduced population of skeletal muscle progenitors. A-D”’. Staining using MET (red) and PAX3 (green) antibodies, and with OAPI (blue) on transverse sections through the fore­ (A-B”’) and hindlimb (C-O”’) bud regions of E11.5 embryos from the compared genotypes (cMet^KO^ and WT). Marked absence of pre-muscle masses in dorsal and ventral portions of the developing fore-(A-A”’) and hindlimb (C-C”’) buds from cMet^KO^ E11.5 embryos compared to those from WT (B-B”’ and 0-0”’) embryos. **E-F.** The number of embryonic muscle progenitor cells co-staining for MET and PAX3 iin the FLBs (E) and HLBs (F) is considerably reduced in cMet^KO^ E11.5 embryos in comparison to the WT embryos. The MET and PAX3 co-positive cell count data were scored using n2:8 non-serial 10µm sections from two embryos of each genotype. The data is represented as means ±SEM and was statistically analysed using a two-tailed unpaired T-test. Scale bar=500µm. D=Dorsal, FLB=Forelimb bud, HLB=Hindlimb bud, and V=Ventral.

Somitic myotome, is the first myogenic structure to form in the developing embryo, while the EMPCs begin migrating from the hypaxial dermomyotome into the developing limb buds^2^. It is therefore expected that somitic myotome adjacent to the limb bud, in WT E11.5 embryos, will show a certain extent of myogenic differentiation. It is important to determine whether disruption of Met signaling in the cMet^KO^ embryos leads to altered myotomal differentiation and morphogenesis, owing to reduced *Slit1* levels seen in the mutant limb buds (Fig. 3B-C and 3F-I). To examine this we first immunostained transverse sections of E11.5 embryos through the limb bud region, for myosin (green), which is a late marker of myogenesis expressed by post-mitotic muscle cells. The nuclei of cells were stained with DAPI (blue). Interestingly, the size of myosin-stained myotome in cMet^KO^ E11.5 embryos (Fig. 7A-A’) was smaller than in WT embryos (Fig. 7B-B’), which was confirmed statistically by analysing somitic myotomal area of three embryos of each genotype and is depicted as a bar graph (Fig. 7C). Next, we stained sagittal sections of E11.5 embryos using anti-myosin antibody and DAPI (blue). We noted that somitic myotome adjacent to FLB was differentiated to a greater extent, comprising long and aligned myofibers, and appeared more compact in cMet^KO^ E11.5 embryos (Fig. 7D-D’) compared to WT embryos (Fig. 7E-E’). Similarly, somitic myotome overlying the HLB also showed a higher degree of differentiation and organization in cMet^KO^ E11.5 embryos (Fig. 7F-F’) than in WT embryos (Fig. 7G-G’). Further, crisp myosin-stained puncta and nascent fiber-like structures marked by white arrows and arrow heads, respectively, are seen distally and proximally in FLB (Fig. 7E) and HLB (Fig. 7G) of WT E11.5 embryos. Whereas, cMet^KO^ E11.5 embryos show mostly diffuse, possibly non-specific, myosin staining distally in fore-(Fig. 7D) and hindlimb buds (Fig. 7F), with a few crisply stained puncta and nascent myofibers restricted to base/proximal region of the limb buds (demarcated by yellow dashed lines in Fig. 7D and 7F-F’). This suggests that EMPCs in WT embryos, which have migrated to the limb buds, have begun terminal myogenic differentiation. Overall, it can be construed that *in vivo* MET signaling is crucial to normal morphogenesis of the somitic myotome close to the limb buds, and in repressing premature myogenic differentiation of migratory EMPCs, ensuring proper migration to their target sites.

**Figure 7.**
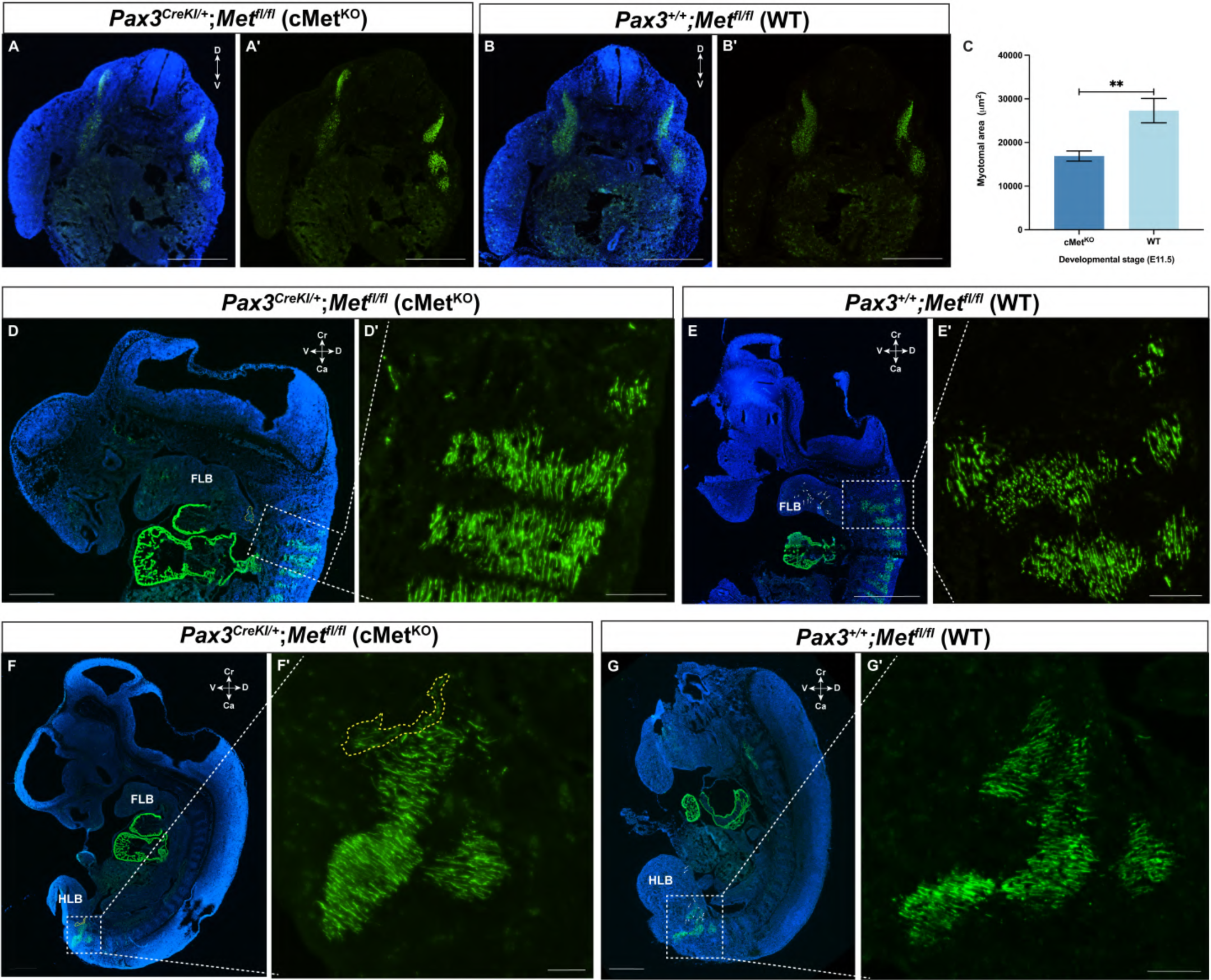
Somitic loss of *Met* alters myotomal size and differentiation. **A-B’.** Representative micrographs showing immunostaining for myosin (green), and nuclei within cells stained with DAPI (blue) in transverse sections through the limb bud region of cMet”^0^ (A-A’) and WT (8-B’) E11.5 embryos, where marked myosin signal is seen in the somitic region. Scale bar=500µm. C. Bar graph depicting substantially reduced myotomal area, quantified from myosin stained sections in cMet”^0^ (A-A’) compared to WT (8-B’) E11.5 embryos (n=3). The graphical data are represented as means ±SEM and the significance was tested using two-tailed unpaired T-test. D-E’. Representative micrographs of DAPI stained sagittal sections from cMet”^0^ (D) and WT (E) E11.5 embryos, and corresponding magnified images of somitic area adjacent to FLB (D’ and E’), marked by white dashed boxes, showing immunostaining for myosin (green). Scale bar is 500µm, 90µm and 80µm in panels D-E, D’ and E’, respectively. **F-G.** Representative sagittal sections of cMet”^0^ (F) and WT (G) E11.5 embryos stained using antibody to myosin (green), and the nuclei were stained with DAPI (blue). **F’-G’.** Magnified images of regions demarcated by white dashed boxes in corresponding panels (F and G), showing myosin staining in somitic myotome overlying the HLB in cMet”^0^(F’) and WT (G’) E11.5 embryos. Scale bar is 500µm, 50µm and 80µm in panels F-G, F’ and G’, respectively. FLB=Forelimb bud, HLB=Hindlimb bud, Ca=Caudal, Cr=Cranial, D=Dorsal, and V=Ventral.

## Discussion

Mammalian skeletal muscle aids voluntary movements, maintains posture and stability, protects visceral organs, allows respiration and ingestion of food, acts as a nutrient reserve, serves as site of metabolism, homeostasis and thermoregulation, and is therefore indispensable for life. Apart from the head and neck all skeletal muscles of the body including those of forelimb and hindlimb (i.e. appendicular muscles), arise from somites^2^. Limb muscles are a valuable model to study myogenesis because they originate from a distinct set of somitic muscle progenitors, further characterized by phase-specific markers that allow conditional mutagenesis to study gene functions crucial to primary and secondary myogenesis. Our study, using mouse genetics identifies *Slit1*, a ligand of the Slit-Robo signaling axis, as a target of MET in signaling deficient cMet^KO^ embryonic limbs. We demonstrate here that MET signaling has an additional non-migratory role in preventing premature myogenic differentiation, which this RTK mediates by regulating *Slit1* expression (Figure 8)

**Figure 8.**
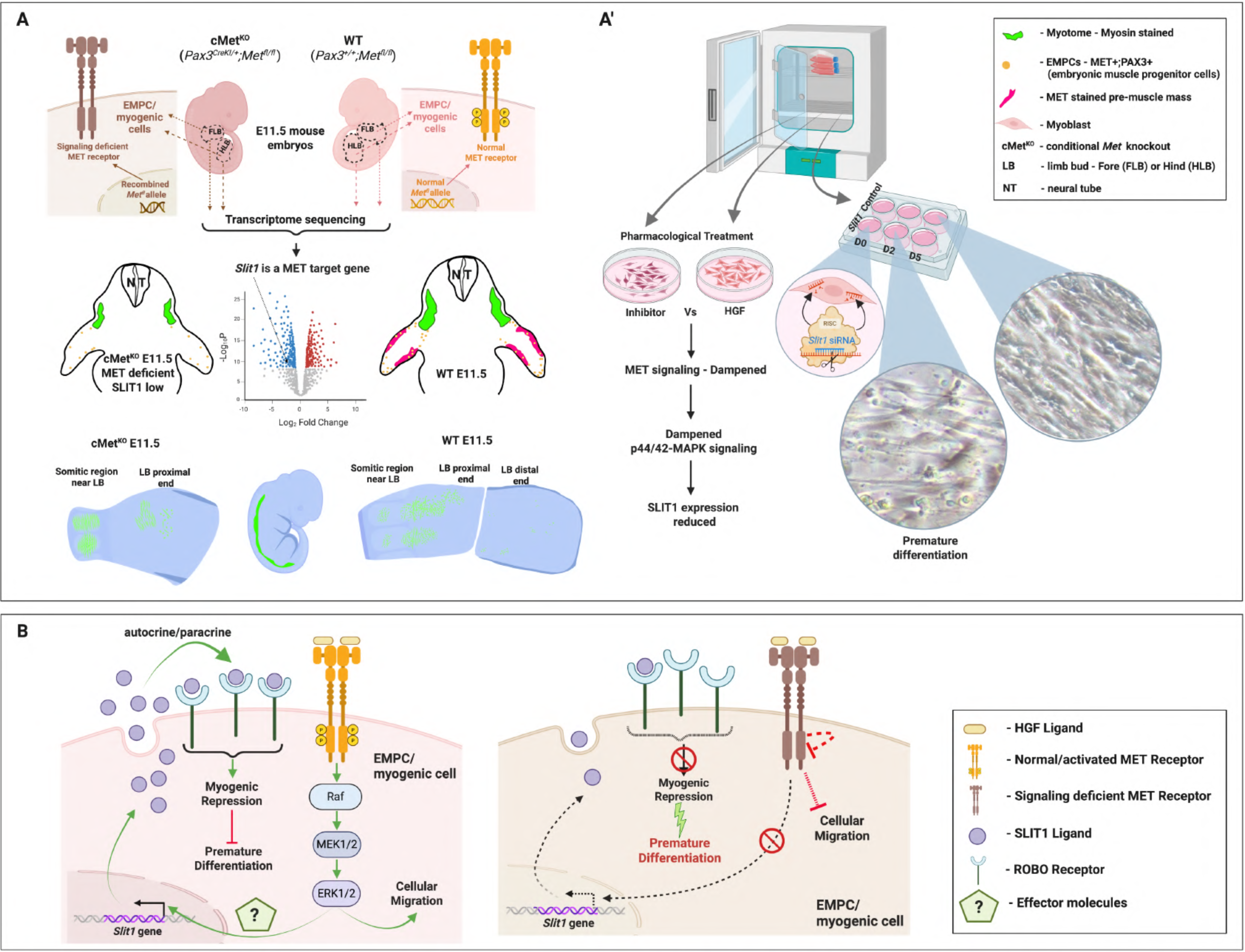
MET signaling fine-tunes mammalian myogenesis through its target *Slit1.* **A-A’,** Genetically ablating *Met* in the somitic/Pax3 lineage (A) or inhibiting the receptor pharmacologically *in vitro* **(A’)** results in reduced *Slit1* expression. Further, downregulating *Slit1, in vitro,* results in premature differentiation of myoblasts which mimics precocious appearance of more differentiated and compactly organised myofibers in appendicular myotomes of cMet^KO^ compared to WT embryos at E11.5. **B_** MET signaling regulates mammalian myogenesis by a two pronged approach - a) causing EMPC migration to the developing limb buds and b) ensuring via SLIT1 that EMPCs remain undifferentiated until they reach the target sites.

Consistent with previous reports^38^, we show that somitic ablation of *Met*, which is expressed in migratory EMPCs, results in muscleless diaphragm in the cMet^KO^ neonates. However, cMet^KO^ fetuses we harvested at E18.5, to score breath counts, appeared pink similar to their non-knockout counterparts, since fetuses rely on maternal blood supply for oxygenation. Presence of cMet^KO^ fetuses at term rules out placental defects in the conditional mutants, that are reported in germline knockout due to deficient MET signaling in trophoblasts^60^. cMet^KO^ fetuses (E18.5) because of amuscular diaphragms develop respiratory distress-induced cyanosis and die within a few hours. Similarly, in the new born litter the transition from fetal to neonatal life involved a shift from reliance on maternal blood supply to active respiration for oxygen intake, which explains the cyanosis observed in the cMet^KO^ neonates (P0). Nevertheless, it remained unclear whether the cMet^KO^ neonate eventually dies because of cyanosis and is then devoured by the mother or long durations of apnea-related lack of spontaneous movement signals the mother to cannibalise the seemingly dead pup.

Further, we noted cMet^KO^ neonates (P0) have highly dysplastic limb muscles unlike amuscular limbs reported in other Met mutants^22,23^. Presence of MET+;PAX3+ EMPCs in limbs of cMet^KO^ E11.5 embryos possibly explains dysplastic appendicular muscles. Though in comparison to WT embryos there was a considerable reduction in EMPC numbers in the cMet^KO^ embryos. Presence of MET+;PAX3+ EMPCs in limb buds of cMet^KO^ embryos suggests that these progenitors are either recombination escapees or there are molecular cues in addition to MET signaling, such as Lbx1, that contribute to their migration^24^. Nonetheless, in the absence of MET signaling, these cues appear inadequate for normal myogenesis. This conclusion is drawn from the observation of loosely and randomly arranged myosin-stained dysplastic muscle cells in cMet^KO^ compared to WT hindlimb sections that show well-formed and organized muscle bundles at P0.

Pax3 is a key myogenic regulator that is expressed in the somite derived EMPCs, which also express Met^24^. *Sp* mice are essentially Pax3 mutants where EMPCs fail to migrate resulting in muscleless limbs^17,18^. In the *Pax3^CreKI^* line used to generate the cMet^KO^ animals, the Cre transgene is knocked-in to replace the first exon of the *Pax3* gene^32^. This results in a functionally null PAX3 protein in the homozygotes leading to an embryonic lethal phenotype^31^. However, in heterozygote animals, Cre mimics the spatio-temporal expression of PAX3, for example Cre is detected in the E9 somites, paralleling PAX3 expression^31^. We investigated the potential impact of hemizygous *Pax3* allele on myogenesis and observed that hindlimbs in *Pax3^CreKI/+^*exhibited normal muscle organization. This precluded any involvement of the hemizygous *Pax3* allele in the dysplastic muscle observed in cMet^KO^ (*Pax3^CreKI/+^;Met^fl/fl^*) neonates. Although limbs from cMet^KO^ neonates (P0) showed dysplastic muscle, innervation (neurofilament staining) was still observed in the limbs. It remains to be determined if the differential muscularization between cMet^KO^ and WT neonates results in changed limb innervation patterns, similar to those seen in the muscleless diaphragm^38^. We noted an apparent increase in expression of *Hgf* (MET ligand) in the cMet^KO^ embryonic limbs compared to WT limb buds. We have not investigated this further, but this may be a temporal compensatory response by the mesenchyme, to inadequate number of EMPCs in the cMet^KO^ embryonic limb buds, of increasing HGF expression to activate more progenitors and promote their delamination and migration.

MET expression is downregulated in the primary myofibers within the limbs at E12.5, but its expression persists distally along the digits, which likely are regions of pre-cartilaginous condensation, and by E14.5, its expression in developing limbs is localized to the future joint regions^26^. We observe remarkably reduced tibio-fibular distance in the limbs of cMet^KO^ neonates (P0) having dysplastic muscles. Given these observations, it may be necessary to a) examine the impact of aberrant appendicular myogenesis on the formation of skeletal structures in the limbs of cMet^KO^ mutants, and b) determine potential direct role of Met in skeletogenesis during embryonic limb formation. This may be relevant particularly due to recent reports associating mutations in the MET kinase domain, which is also the region recombined out in the cMet^KO^ mutants, to human familial distal arthrogryposis and muscular dysplasia^27,28^.

We also identified *Slit1*, among other candidates, as a differentially expressed gene in the transcriptome datasets from RNA sequencing of fore- and hindlimb buds of cMet^KO^ and WT E11.5 embryos (Fig. 3B and 3C). The Slit1-Robo2 signaling axis is involved in chick myogenesis^54^, and *Slit3* knockout mice were shown to develop congenital diaphragmatic hernia (CDH)^61^. Our data and these reports suggest that *Slit* family members have roles in appendicular and diaphragmatic myogenesis. SLIT ligands and ROBO receptors are encoded by families of homologous genes. Given *Slit1* downregulation in cMet^KO^ embryonic limbs, we checked for compensatory increase in its homologs’ expression. We also tested if *Robo2*, which encodes the cognate receptor of SLIT1 ligand, and its homologs were altered in cMet^KO^ limb buds. We quantified levels of *Slit* and *Robo* homologs by qPCR, and expression of representative homologs in FLBs and HLBs is shown in Fig. 3H and 3I, respectively. Only *Slit1* shows substantially decreased expression in cMet^KO^ compared to WT embryonic limbs, suggesting that the recombined *Met* allele in somite derived EMPCs produced a signaling deficient protein incapable of maintaining normal *Slit1* expression (i.e. cells were *Slit1* low). Thus, *Slit1* appears to be a target of MET signaling in migratory progenitors crucial to embryonic appendicular myogenesis. Further, the MET signaling deficient-*Slit1* low EMPCs with impaired migratory potential exhibited a more differentiated and aligned/ordered myogenic structure in the myotome adjacent to the limb buds in cMet^KO^ embryos compared to WT E11.5 embryos. This observation is reinforced by the reduced myotomal area or more compact myotome (myosin stained) seen in sections through the limb bud region in cMet^KO^ E10.5 embryos. Likewise, in vitro, *Slit1* expression changed in accordance with pharmacological inhibition of MET signaling, and knocking down its levels in myoblasts caused them to fuse precociously to form myotubes at earlier timepoints in differentiation. Thus, through genetic manipulation and pharmacological inhibition, we demonstrate that, beyond regulating EMPC migration, MET has a function in preventing premature progenitor differentiation during mammalian myogenesis, which is mediated by *Slit1*. Given that skeletal muscle formation is crucial to survival and quality of life postnatally, it is important to examine Met-mediated spatio-temporal regulation of *Slit1* and to explore whether other independent cues converge to execute the concerted functions of *Met* and *Slit1* in mammalian myogenesis.

## Materials and Methods

### Mice lines, developmental staging and PCR genotyping

The *c-met^fl^* is a conditional allele denoted as (*Met^fl^*), and the Cre driver line - *Pax3^CreKI^* is a knock-in mutant mouse strain^29,32^. Both these lines -*Met^fl^*(RRID:IMSR_JAX:016974) and *Pax3^CreKI^* (RRID:IMSR_JAX:005549) were obtained from The Jackson Laboratory, Bar Harbor, Maine, USA^30,31^. Timed matings were set-up to harvest embryos at specific developmental stages, considering the morning of vaginal plug in the dams as embryonic day 0.5 (E0.5). Wildtype (WT) and knockout (cMet^KO^) embryos were collected in PBS over ice from crossings *Pax3^CreKI^; Met^fl/+^* and Met^fl/fl^ animals. Animal maintenance and all experiments were carried out in accordance with protocols approved by the Regional Centre for Biotechnology institutional animal ethics committee (IAEC protocol numbers: RCB/IAEC/2017/020 and RCB/IAEC/2024/204).

The Wildtype, Lox and Cre alleles were identified using conventional PCR genotyping. Briefly, ear clips from adult mice and tail clips from embryos and neonates were used to extract genomic DNA by the HotSHOT lysis method (Truett Warman Bio Techniques 2000)^62^. 15μl genotyping PCR reaction comprised 7.5μl GoTaq G2 Hot Start Green Master Mix (Promega, Cat # M7423), 0.75μl each of 10μM forward and reverse primers (0.5μM final concentration), 5μl HotSHOT DNA and 1μl nuclease free water. Primer sequences and PCR cycling conditions are detailed in Supplementary Table 1. PCR products were analysed by electrophoresis on 1.5% agarose gel in 1x TAE (Tris Acetate EDTA) buffer, and alleles were attributed based on band sizes detailed in Supplementary Table 1.

### Whole-mount immunohistochemistry

Pups collected at post-natal day 0 (P0) were euthanized by decapitation following isoflurane-induced anesthesia. Tail snips were used for genotyping. Diaphragm explants were prepared by cutting the P0 through the thoracic and abdominal cavities indicated along the lines 1 and 2, respectively (Supplementary Fig. 1A). Thus, the body of the P0 neonate was dissected into 3 portions – a) torso with the diaphragm, b) head portion with forelimbs and c) caudal portion with the hindlimbs, which were processed for whole-mount immunohistochemistry (WM-IHC) with slight modifications to the protocols described previously^38,63,64^. Briefly, the P0 explants were fixed in 4% PFA at 4°C on a rocker (R) for 24 hours (h), rinsed with 1x Phosphate buffered saline (PBS) and then incubated in Dent’s bleach at 4°C/R/24-h. Embryos were rinsed with 100% methanol and fixed in Dent’s fixative at 4°C/R/5 days.

After fixation, the heart, lungs, liver and gut were removed carefully from the torso, so as not to damage the diaphragm. Limb explants were removed from the head and caudal portions of the P0 and were deskinned in 100% methanol prior to IHC staining. The epimysium covering the muscle was removed very carefully. The limb and diaphragm explants were rehydrated in 50% methanol (in 1x PBS) at room temperature (RT)/R/10 minutes (min).

Prior to labeling with Neurofilament (NF) antibody, the explants were washed thrice, in TBST (1x TBS + 0.1% Tween) at RT/R/5 min, each time. Limb and diaphragm explants were initially bleached and then blocked at RT/R/1-h each, in 6% H_2_O_2_ (in 0.1% TBST) and in blocking solution (5% goat serum + 20% DMSO + 75% TBS), respectively.

Washing and bleaching steps for explants that were labeled with alkaline phosphatase (AP)-conjugated anti-Myosin primary antibody, were performed in PBST (PBS + 0.1% Tween). An additional step was performed after bleaching and prior to blocking, where explants were incubated in PBS at 65°C on a thermo mixer (Eppendorf) for an hour to heat inactivate the endogenous AP activity. After blocking (5% goat serum + 20% DMSO + 75% PBS) at RT/R/1-h, explants were incubated in primary antibody (prepared in the respective blocking solution as per Supplementary Table 3), at 4°C/R/48-h. The explants were cleared in BA:BB solution (33% benzyl alcohol + 66% benzyl benzoate + 1% TBS) at RT/5 min and then rinsed extensively in PBS with each wash lasting 5 min, until the colloidal appearance of the PBS was not seen. Detection was performed by incubating explants with NBT-BCIP premix (Merck, Cat# B6404) diluted to 75% in TBS, with intermittent washes in water to check staining intensity under the dissection microscope. Finally, color development was stopped by rinsing in 1% acetic acid solution twice for 2 min.

Explants stained for NF were incubated with horse radish peroxidase (HRP)-conjugated anti-rabbit secondary antibody provided in the HRP/DAB IHC Detection Kit (Abcam Cat# ab236466), at 4°C/R/24-h. Prior to, and after staining with the HRP-conjugated secondary antibody, the explants were washed thrice in TBST for 5 min, twice in TBST for 30 min and thrice in TBS for 5 min. Each repeat wash was performed at RT on a rocker in the specified solution for the indicated duration. These explants were cleared in a similar manner to explants stained for AP-Myosin, HRP detection was performed as per the manufacturer’s specifications.

Finally, the stained explants were rinsed in TBS, post-fixed in 4% PFA + 0.2% glutaraldehyde at RT/R/10 min, mounted in DPX (Merck; Cat# 06522) and were imaged using the Leica S8 APO and Nikon SMZ 745T microscopes equipped with the LAS X and NIS-Elements softwares, respectively.

### Immunofluorescence

E11.5 embryos (*Pax3^CreKI/+^; Met^fl/fl^* and *Pax3^+/+^; Met^fl/fl^*) were harvested from timed matings (described above in the section on mouse lines). The embryos were processed for staining. Briefly, embryos were extensively rinsed in PBS, fixed in 4% paraformaldehyde (PFA) at RT/R/1-h, rinsed in PBS, equilibrated in a sucrose gradient by immersing initially in 10% sucrose in PBS at 4°C/R/overnight and then in 20% sucrose in PBS at RT/R (incubation in sucrose is continued till the sample sinks) and embedded in optimal cutting temperature medium (OCT -PolyFreeze, Polysciences; Cat# 19636). Hindlimbs of *Pax3^CreKI/+^; Met^fl/fl^* and *Pax3^+/+^; Met^fl/fl^* neonates at P0 were dissected and embedded in OCT and flash-frozen in 2-methyl butane (Sigma; Cat# M32631) pre-cooled using liquid N_2_. OCT-embedded tissues were sectioned to 10μm thickness using a cryostat (SLEE MNT), and the adjacent sections were collected on a series of appropriately labelled coated slides (VWR; Cat# 631-0108). Embryos were sectioned in either transverse or sagittal planes depending on the experimental requirement. Transverse sections close to the middle of P0 limb were used for staining.

For immunofluorescence staining sections of E11.5 embryos and P0 limbs were fixed in PFA at RT/5 min. Antigen retrieval is recommended for the antibodies used in immunofluorescence and was performed by immersing slides in citrate buffer (which comprised 1.8 mM citric acid and 8.2 mM sodium citrate in water) and heating in an antigen retrieval chamber. Tissue sections were cooled to RT, P0 tissues were blocked with 5% goat serum (HiMedia; Cat# RM10701) in PBS, and E11.5 tissues were blocked in 2.5% GS in PBS containing 0.1% Triton-X-100 (MP Biochemicals; Cat# 194854), at RT/1-h. For staining E11.5 tissues with anti-Met antibody, which is raised in goat, 2.5% horse serum (Thermo Fisher Scientific; Cat# 16050122) in PBS containing 0.1% Triton-X-100 was used for blocking. The samples were incubated in optimized concentration of primary antibody (Supplementary Table 3) overnight at 4°C in a moist chamber, incubated with appropriate secondary antibody at RT/2-h in a moist chamber. Each step was followed by rinsing thrice in PBS at RT/5 min. Samples were post-fixed in 4% PFA at RT/5 min, rinsed in distilled water, air dried and mounted using DAPI Fluoromount-G (Southern Biotech; Cat# 0100-20). Fluorescence microscopy was performed using a Nikon Eclipse Ti2 microscope. ImageJ software was used for counting stained cells or measuring size of stained regions.

### RNA isolation, transcriptome sequencing and analysis

A pair of forelimb buds (FLBs) was harvested from four conditional Met knockout (cMet^KO^) and four wildtype (WT) mouse embryos at E11.5. Similarly, a pair of hindlimb buds (HLBs) was harvested from four E11.5 embryos of each genotype (cMet^KO^ and WT). RNA was extracted using RNeasy Lipid Tissue Mini Kit (Qiagen; Cat# 74804). Briefly, QIAzol lysis reagent was added to the tissue in the microcentrifuge tube, the sample was macerated using a plastic pestle and RNA was isolated according to the manufacturer’s protocol. Integrity and quality of the isolated RNA was verified using the RNA 6000 Nano Kit (Agilent; Part no. 5067-1511) on the 2100 Bioanalyzer (Agilent; Part no. G2938C) following the manufacturer’s specifications.

1μg aliquots of RNA samples having RIN in the range of 8.1 to 9.9 were shipped to the Genomics Core Facility (GCF), Washington State University, Spokane, for RNA-sequencing. At GCF, the integrity and quality of samples was re-assessed using a 5200 Fragment Analyzer System (Agilent; Part no. M5310AA) and concentration measured on a Qubit^®^ 2.0 Fluorometer (Thermo Fisher Scientific; Part no. Q32866). RNA samples were of high quality and library preparation was performed using Illumina TruSeq stranded mRNA library preparation kit (Illumina; Cat# 20020594) with oligo-dT beads to selectively capture polyadenylated transcripts, as per the manufacturer’s protocol. Paired end 151bp reads were generated by sequencing the libraries on the Novaseq X Plus system (Illumina; Cat# 20084804).

Read quality was assessed by FastQC, and low-quality reads were trimmed using Trimmomatic. While the TruSeq mRNA Library Prep kit, through its poly-A enrichment approach, enriches for mRNA and inherently depletes most rRNA, SortMetRNA software was used to filter rRNA fragments from meta-transcriptomic data. This additional precaution ensures minimal rRNA contamination in the final dataset. Reads were mapped to the mouse reference genome [Mus musculus, GRCm38 (Ensembl release 102)] using HISAT2.

Read counts were assessed by featureCounts and differential expression analysis was performed using the DESeq2 package in R. Differential gene expression was evaluated using a cutoff P < 0.05, and | log2(Fold Change) | ≥ 1, or ≤ −1, which signifies at least 2-fold upregulated and downregulated gene expression, respectively. A Venn diagram was made using Venny2.1.0 and bioRender, to identify DEGs that are common between forelimb and hindlimb buds and unique to specific developing appendages in cMet^KO^ and WT embryos (E11.5). Gene ontology (GO) enrichment analysis for biological processes was performed with DEGs that were significantly regulated (based on the above-mentioned filtering parameters) using ShinyGO 0.82 software.

To further assess the RNAseq data quality, graphical visualizations such as dispersion plots, principal component analysis (PCA) plots and volcano plots were generated using the filtered and normalized datasets of FLBs and HLBs from the two compared groups, using DESeq2 and EnhancedVolcano packages in RStudio. The QC analysis using dispersion and PCA plots is presented in Supplementary Fig. 2. Heatmaps were prepared using the ComplexHeatmap package in RStudio^65^ and the criteria to filter DEGs between compared groups were padj < 0.05 and | log2(Fold Change) | ≥ 1.5, or ≤ −1.5, i.e. at least 3-fold change in expression.

### cDNA synthesis and quantitative PCR

Expression levels of specific candidate genes identified by RNAseq were validated by quantitative real-time polymerase chain reaction (qPCR), employing the same RNA samples that were used for RNAseq. cDNA was synthesized from approximately 1μg RNA using High-Capacity cDNA Reverse Transcription Kit (Applied Biosystems, Thermo Fisher Scientific; cat# 4368814).

Primer3, an online software^66^, was used to design primers specific to candidate genes/ genes of interest (GIs – *-Met*, *Hgf*, *Cdh15/M-Cad*, *Msc/MyoR*, *Slit1*, *Slit3*, *Robo2* and *Robo4*) and reference genes (GRs – *Gapdh, Ap3d1 and Ppia*). Specificity and efficiency of the primers were tested as detailed previously^67^. Since the *Met^fl^*allele carries a floxed exon 16 that gets recombined out and deleted in the progeny, by crossing with the *Pax3^CreKI^* driver line. The cMet^KO^ will therefore have a truncated MET protein that is devoid of only exon 16. Accordingly, qPCR primers were designed specifically to the exon 16 of the *Met* gene to validate the knockout. Details of primers are provided in Supplementary Table 2.

Transcripts were quantified using SYBR Green (Applied Biosystems, Thermo Fisher Scientific; Cat# A25742) on the QuantStudio™ 6 Flex Real-Time PCR System (Applied Biosystems, Thermo Fisher Scientific; Cat# 4485691). All the assays were set-up as described previously^67^. qPCR data were analysed on the QuantStudio™ Real Time PCR Software v1.3 (Applied Biosystems, Thermo Fisher Scientific). Expression levels of GIs in the cMet^KO^ samples relative to WT samples, were calculated using the relative quantitation (RQ = 2^−ΔΔCt^, Ct=threshold cycle) method^68^. Briefly, the expression of GIs was first normalized to the expression of GRs in the respective samples (ΔCt = Avg Ct GI – Avg Ct GR, Avg Ct = average cycle threshold value calculated from concordant triplicate Ct values). The mean expression value of two GRs was used for normalizing GIs, and the choice of the GRs was guided by their stable expression across cMet^KO^ and WT samples in FLBs (*Gapdh* and *Ap3d1*) and HLBs (*Gapdh* and *Ppia*), respectively. In literature, *Gapdh*, *Ap3d1* and *Ppia* have been used as reference for normalizing target gene expression in murine myoblast C2C12 cells, developing mouse forelimb and human myoblasts, respectively^69,70^. The normalized GI expression of the cMet^KO^ and WT samples was in turn used for a second round of normalization (ΔΔCt = ΔCt cMet^KO^ - ΔCt WT) to compute the relative expression value of GIs in the cMet^KO^ samples. Four biological replicates were used for both FLBs and HLBs.

### Cell culture

C2C12 mouse myoblast cells (American Type Culture Collection-ATCC; Cat# CRL-1722) were cultured and maintained in complete medium [CM = DMEM (Thermo Fisher Scientific; GIBCO, Cat# 11995065) + 10% fetal bovine serum (FBS, Thermo Fisher Scientific; GIBCO, Cat# 10270106) + 1% Penicillin-Streptomycin (Thermo Fisher Scientific; GIBCO, Cat# 15240062)], according to ATCC recommendations.

Pharmacological treatment was carried out in C2C12 cells to study the effects of specific reagents on Met signaling and its downstream cascades in modulating the expression of genes identified from the RNAseq screen. Briefly, proliferating C2C12 myoblasts were seeded at a cell density of 1.52x 10^5^ cells per well, of a 6 well cell culture plate, in CM. Cells were allowed to proliferate for 24-h and attain ∼80% confluence, following which they were treated with optimized concentrations of Met ligand i.e. HGF (Thermo Fisher Scientific, Peprotech; Cat# 315-23), Met inhibitor SU11274 (Cayman Chemical, Cat# 14861), MEK1/2 inhibitor U0126 (Cell Signaling Technology, Cat# 9903S) and Wortmannin (WO, Cayman Chemical, Cat#10010591) at final concentrations of 10ng/ml, 25nM, 20μM and 2.5μM, respectively, for durations indicated below. HGF was solubilized in 0.1% BSA (prepared in sterile 1x PBS), whereas SU11274, U0126 and WO were solubilized in DMSO.

C2C12 cells were also treated also with the solvents/vehicles used to solubilize the respective ligand and inhibitors, at concentrations matching their final concentrations in wells subjected to corresponding pharmacological treatment. Lysates of C2C12 cells treated correspondingly with the solvents/vehicles were used as controls/baseline expression of target genes to normalize their expression upon pharmacological treatment.

HGF-24-h, SU11274 – 24-h, and SU11274 – 24-h, followed by HGF – 2.5-h treatment prior to harvesting (TPH) HGF-2.5-h, U0126 – 2.5-h, and U0126 – 2.5-h, followed by HGF – 30 min TPH HGF-2.5-h, WO – 2.5-h, and WO – 2.5-h, followed by HGF – 30 min TPH Gene knockdown was performed by reverse transfecting target and control siRNAs in C2C12 cells that were subsequently subjected to a differentiation protocol. Briefly, 3×10^4^ C2C12 cells suspended in 400μl CM were plated per well of a 24-well cell culture plate containing the transfection mixture. Transfection mixture per well comprised 100μl Opti-MEM (Thermo Fisher Scientific; GIBCO, Cat# 31985070), 50nM silencer select™ pre-designed *Slit1* (Thermo Fisher Scientific; Ambion, Cat# 4390771, ID.: s73981) or negative control siRNA no. 2 (Thermo Fisher Scientific; Ambion, Cat# 4390846) and 2μl Lipofectamine RNAiMAX transfection reagent (Thermo Fisher Scientific; Cat# 13778150). Cells were grown in this CM + transfection mix for 48-h to allow growth to approximately 80% confluence and ensure efficient transfection. Cells were harvested from at least 3 replicate wells 24-h after transfection, cell extracts were prepared as described below in the western blotting section and assigned as differentiation day zero (D0) lysates (Supplementary Fig. 1C). 24-h later differentiation was induced in the remaining wells by replacing CM + transfection mix with differentiation medium (DM = DMEM + 2% horse serum + 1% Penicillin-Streptomycin) and then replaced 50% spent DM with fresh DM every 24-h for the next 144 hours. Cells were harvested and lysates prepared on specific days (D) of the myogenic differentiation protocol indicated in Supplementary Fig. 1C.

### Western blotting

Protein extracts were prepared from C2C12 cells that were plated, cultured and harvested at specific time points as described above in the section on cell culture. Briefly the spent medium was aspirated from the wells and the cells were trypsinized (GIBCO Cat# 25200056). Trypsin was neutralized with complete medium, the cell suspension was aspirated into a Falcon tube and centrifuged at 6000 rpm/4°C/6 min. Cells were washed twice in ice-cold 1x DPBS (Gibco; Cat# 14190-144) and centrifuged at 6000 rpm/4°C/6 min each time. Cells were lysed in ice-cold radio-immunoprecipitation assay (RIPA) buffer (Sigma; Cat# R0278) supplemented with Protease/Phosphatase Inhibitor Cocktail (100X, Cell Signaling Technology Cat# 5872S) at 1:100 dilution. The protein in the lysates was quantified using Pierce BCA Protein Assay Kit (Thermo Fisher Scientific; Cat# 23225) as per supplier’s protocol. 7% or 10% SDS-PAGE gels were used, depending on the molecular weight of the protein to be detected, to separate equal concentrations of protein extracts. Subsequently, the electrophoresed proteins were transferred to PVDF membrane (Millipore; Cat# IPVH00010), the membranes were blocked in 5% skimmed milk or 5% Bovine serum albumin (BSA) or skimmed 3% milk:2% BSA for 5-h (as optimised for specific antibody), incubated overnight in primary antibody at 4°C and 2-h in an appropriate HRP-conjugated secondary antibody at RT. Protein signal was detected using Clarity Western ECL Substrate (Biorad; Cat# 1705060) and blots were imaged on the ImageQuant LAS 4000. Densitometric analyses were performed using the ImageQuant software or ImageJ software version 1.54p^71^. Reference genes (GRs) used as loading controls for normalizing protein expression were β-ACTIN, GAPDH, or ⍺-TUBULIN. Data from at least three independent replicates were plotted as an average, with the error bars representing ± standard deviation (SD). In case of pharmacological treatments, treatment-responsive protein expression was normalized first to GR controls and then to the corresponding solvent/vehicle treatment, which had already been normalized with their respective GR controls. Protein expression of the genes of interest (GIs) at specific day of myogenic differentiation (D2, D5 & D7), in control and *Slit1* siRNA transfected C2C12 cells, was normalized first to the corresponding reference gene/loading control (i.e. GI/GR), which was in turn normalized to the D0 (GI/GR) value of respective transfection and plotted adjacent. Antibodies used for western blotting are listed in Supplementary Table 3.

### Schematics, Graphing and Statistical analyses

The graphical abstract (Fig. 8) and schematics (Fig. 1A, 3A and Supplementary Fig. 1A) were prepared using bioRender. Experimental data were plotted and statistically analysed in GraphPad Prism Version 10.5.0 f(697) (GraphPad Software Inc., CA, USA). Bar graphs summarising the western blot densitometry data are represented as means ±SD. All other graphical data is represented as means ±standard error of mean (SEM), and each experiment was performed at least in triplicate. Statistical tests that the data were subjected to, are specified in the figure legends. The P-value is indicated on the graph along with asterisks and p-value≤0.05 is considered significant. The level of significance is indicated as *P < 0.05; **P < 0.01; ***P < 0.001; ****P < 0.0001.

## Acknowledgements

We acknowledge valuable suggestions and help from Dr. Perumal Nagarajan, Experimental Animal Facility, National Institute of Immunology, New Delhi, and Dr. Aurile Jory, InStem, Bangalore, with animal work. We also thank Dr. Perumal Nagarajan for critical inputs on the manuscript. MS acknowledges valuable suggestions and inputs from Prof. Shubha Tole and Dr. Sandhya P. Koushika, Tata Institute of Fundamental Research, Mumbai. We thank Dr. Liu Yiyong and Dr. Wang Archer of the Genomics Core Facility, Washington State University, Spokane, for helping with RNA sequencing. Staff of the Small Animal Facility (SAF) at the NCR Biotech Cluster, and Ms. Diksha Singh, staff on the grant/funding to MS, are duly acknowledged for their assistance with animal work. We also acknowledge staff of the central instrumentation and microscopy facilities at the Regional Center for Biotechnology (RCB), Faridabad, for their help. MS and JJ thank Mr. Naveen Kumar for technical help. We acknowledge the support of DBT e-Library Consortium (DeLCON) for providing access to e-resources.

## Data availability

RNA-Seq datasets generated in this study have been deposited in Indian Biological Data Centre (IBDC) and International Nucleotide Sequence Database Collection (INSDC) under the accession numbers INRP000194 and PRJEB82980, respectively.

## Author contributions

MS conceptualized the study, acquired funding, designed and performed the experiments, curated, analysed and interpreted the data, and wrote the manuscript. JJ assisted with animal maintenance, performed experiments, curated, analysed and interpreted data. GC guided MS in experimental design, data interpretation, and provided overall supervision. SJM was the local host and mentor for MS and JJ for the entire duration of the project. All authors read, edited and approved the final manuscript.

