## Supplementary Figure 1 for "*Slit1* -a MET target gene in the embryonic limbs, prevents premature differentiation during mammalian myogenesis"

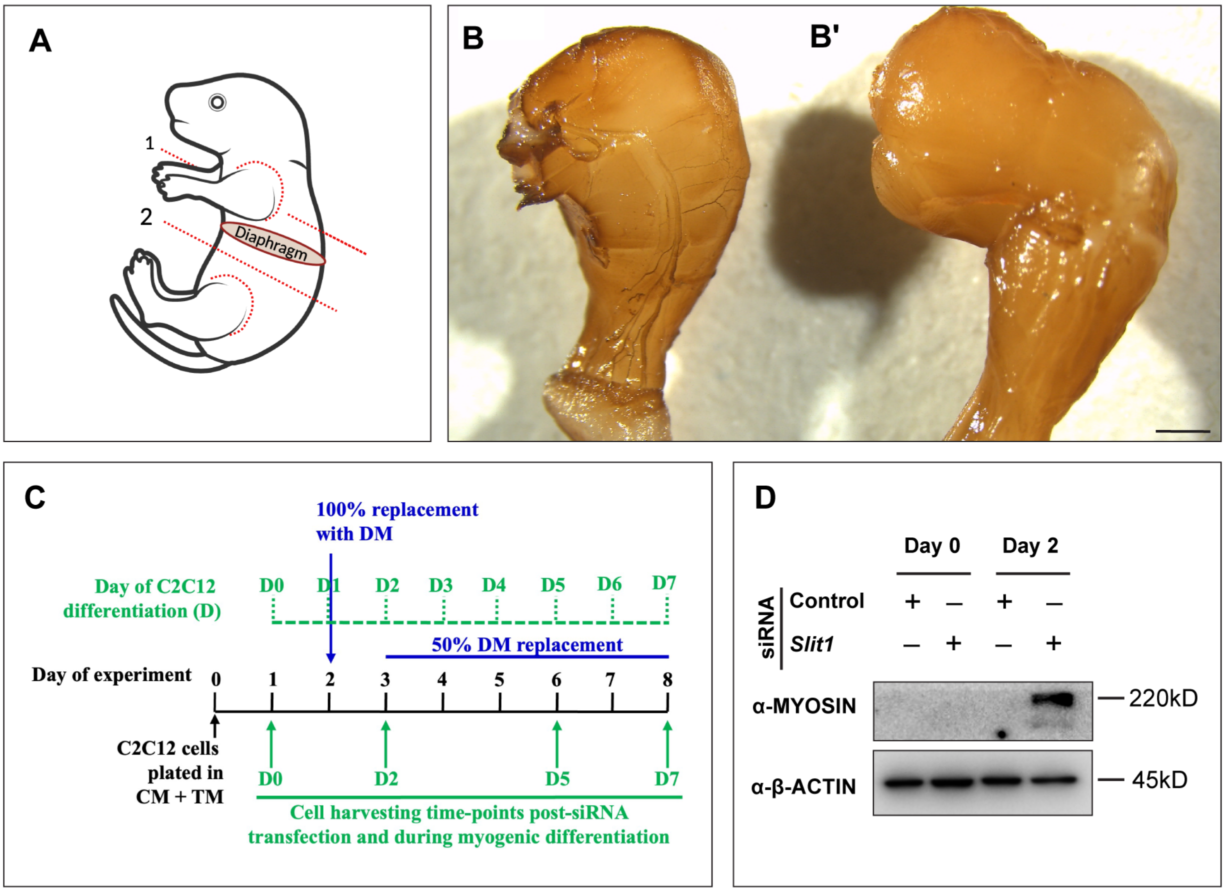


**Supplementary Figure 1. Generating organ explants from P0 neonates for IHC, testing specificity of neurofilament antibody, inducing differentiation and examining the effect of *Slit1* knockdown in mouse myoblasts. A.** Schematic representing dissection strategy adopted to generate forelimb, diaphragm and hindlimb explants from postnatal day 0 (P0) mouse pups, for whole-mount immunohistochemistry. **B** and **B'**. Whole-mount of right (B) and left (B') hindlimbs of a WT neonate (P0), incubated with and without primary neurofilament antibody, respectively, and stained with chromogenic substrate (DAB). Scale bar=1000µm, DAB=diaminobenzidine. **C.** Schematic summarising protocol followed for siRNA mediated knockdown in C2C12 cells prior to inducing their myogenic differentiation, and time-points where cells were harvested to prepare lysates for immunoblotting. **D.** Representative immunoblots showing levels of MYOSIN and β-ACTIN in cell lysates of C2C12 cells transfected with control or *Slit1* siRNA, at D0 and D2 of myoblast differentiation. CM=complete medium, TM=transfection mixture, DM=differentiation medium.
