## Supplementary Figure 2 for "*Slit1* -a MET target gene in the embryonic limbs, prevents premature differentiation during mammalian myogenesis"

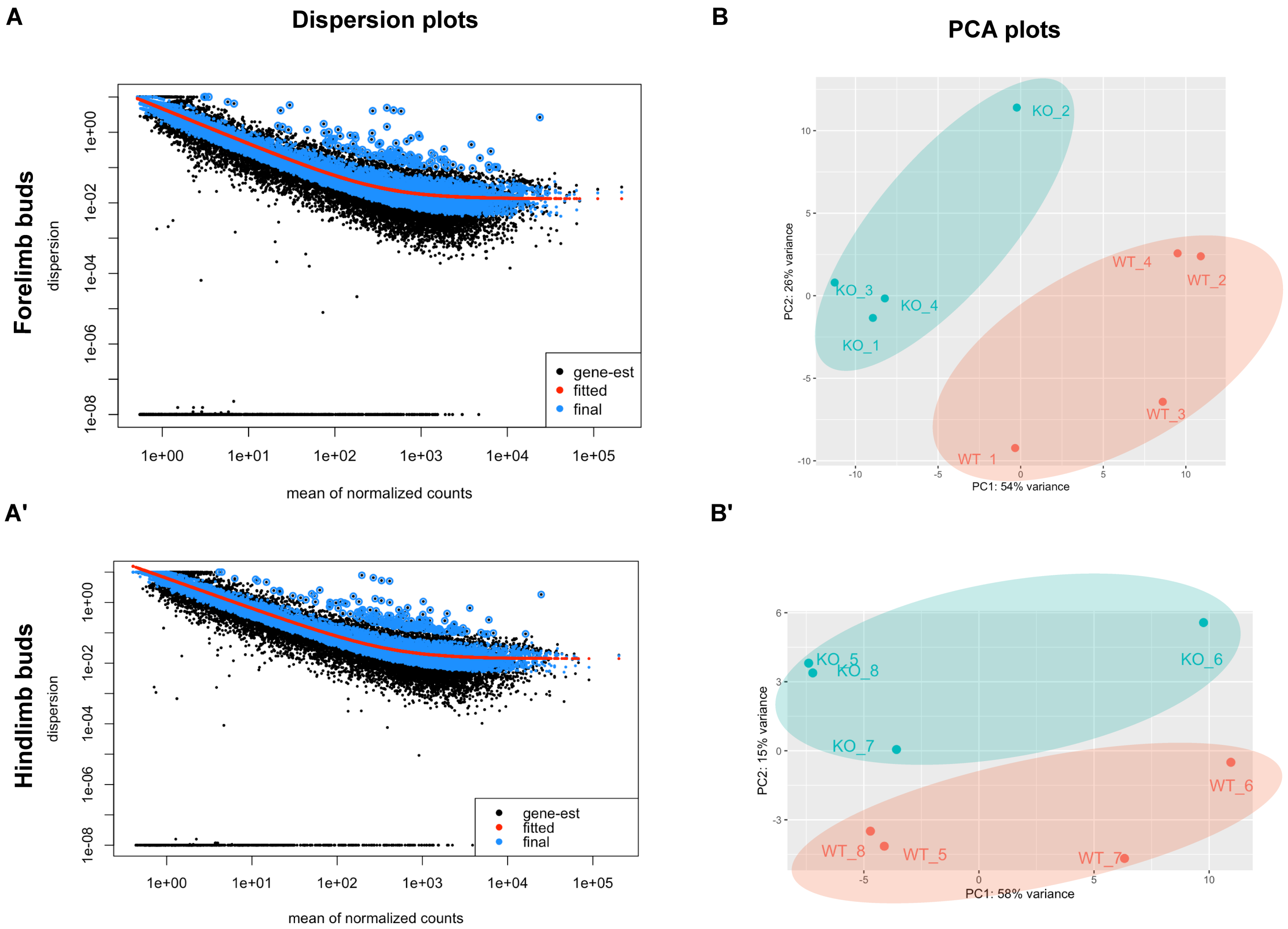


**Supplementary Figure 2. Transcriptome analyses of limb buds between wildtype and conditional *Met* knockout embryos at E11.5. A, A'.** Dispersion plots representing variation in mean expression levels of genes across FLBs (A) and HLBs (A') of conditional *Met* knockout (cMet$^{KO}$) and wildtype (WT) embryos (E11.5). Each black dot represents dispersion of a particular gene. **B, B'.** Principal component analysis (PCA) plots show segregation of compared samples i.e. KO=cMet$^{KO}$ and WT E11.5 embryos, based on variability in transcriptome-wide expression profile between sequencing replicates of FLBs (C) and HLBs (C'). Percentage variation captured by each principal component is represented on the respective axis. FLBs=Forelimb buds, HLBs=Hindlimb buds.
