## Supplementary Figure 3 for "*Slit1* -a MET target gene in the embryonic limbs, prevents premature differentiation during mammalian myogenesis"

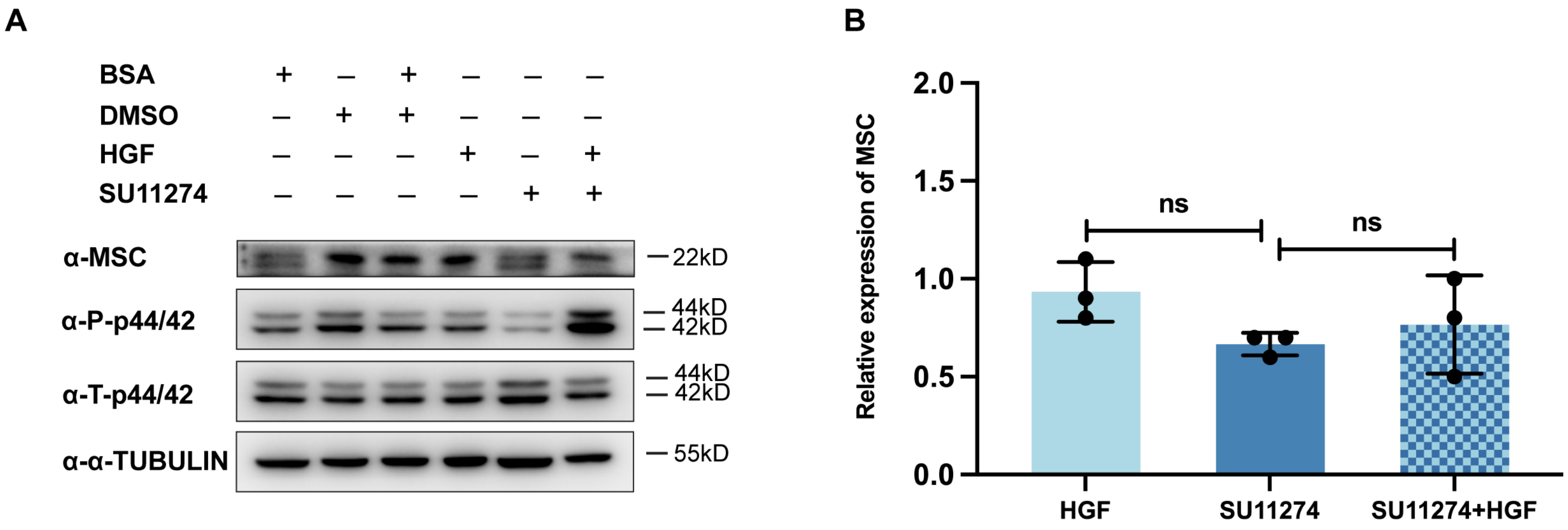


**Supplementary Figure 3. Musculin expression varies in a MET signaling responsive manner. A.** Representative immunoblot showing levels of Musculin (MSC), phosphorylated (P)-p44/42, total (T)-p44/42 and α-TUBULIN in lysates of C2C12 mouse myoblasts in response to MET signaling activation by HGF, inhibition with SU11274, sequential treatment with SU11274 and HGF, and treatment with respective solvents/vehicles, which were used to solubilize these pharmacological reagents. **B.** Bar graph depicting densitometric quantitation of MSC expression determined by western blot (A), normalized first to α-TUBULIN and then to MSC levels (normalized by reference gene) in the respective vehicle controls. Densitometry data are represented as means ±SD and were analysed using one-way ANOVA statistical test.
