## Supplementary Information for "*Slit1* -a MET target gene in the embryonic limbs, prevents premature differentiation during mammalian myogenesis"

**Supplementary Table 1. Primers and PCR cycling conditions used for genotyping**

<table>
<tr><th>S. No.</th><th>Allele symbol</th><th>Primer No.</th><th>Primer sequence (5' → 3') and length (bp)</th><th>Amplicon size (bp)</th><th>Reference/ designed de novo (DD)</th></tr>
<tr><td rowspan="3">1</td><td rowspan="3">Pax3<sup>CreKI</sup></td><td>MS_1115</td><td>FP: ATT GCT GTC ACT TGG TCG TGG C (22)</td><td rowspan="2">Cre mutant – 200<br>Cre wildtype – no band</td><td rowspan="2">Ishan Liu<br>Dev. 2023</td></tr>
<tr><td>MS_1116</td><td>RP: GGA AAA TGC TTC TGT CCG TTT GC (23)</td></tr>
<tr><td>Cycling conditions</td><td colspan="3">Initial denaturation (95˚C, 2 minutes),<br>35 cycles of<br>denaturation (94˚C, 20 seconds),<br>annealing 59˚C, 30 seconds),<br>extension (72˚C, 30 seconds), followed by<br>an additional extension (72˚C, 10 minutes), and<br>holding (4˚C, 30 minutes)</td></tr>
<tr><td rowspan="3">2</td><td rowspan="3">Met<sup>fl</sup></td><td>MS_1478</td><td>FP: TGA CTC ATT TTG CCC AGA GA (20)</td><td rowspan="2">Met<sup>fl/fl</sup> – 600<br>Met<sup>fl/+</sup> – 600 & 470<br>Met<sup>+/+</sup> – 470</td><td rowspan="2">DD</td></tr>
<tr><td>MS_1479</td><td>RP: TTC CAG GTG GCT TCA AAT TC (20)</td></tr>
<tr><td>Cycling conditions</td><td colspan="3">Initial denaturation (95˚C, 3 minutes),<br>35 cycles of<br>denaturation (94˚C, 1 minute),<br>annealing 56˚C, 40 seconds)<br>extension (72˚C, 1 minute), followed by<br>an additional extension (72˚C, 5 minutes), and<br>holding (4˚C, 30 minutes)</td></tr>
</table>

**Supplementary Table 2. List of primers used for quantitative PCR**

| S. No. | Gene symbol | Primer No. | Primer sequence (5' → 3') and length (bp) | Amplicon size (bp) | Reference/ designed de novo (DD) |
|---|---|---|---|---|---|
| 1 | *Gapdh* | MS_646 | FP: GAC TTC AAC AGC AAC TCC CAC T (21) | 169 | Kumar Mathew The FASEB J. 2023 |
| | | MS_647 | RP: GGT CCA GGG TTT CTT ACT CC (20) | | |
| 2 | *Ap3d1* | MS_125 | FP: TGT CTG CAA GCT CAC CTA CT (20) | 153 | DD |
| | | MS_126 | RP: ATG ACG TCG GTA CCT TCA TG (20) | | |
| 3 | *Ppia* | MS_113 | FP: ACC AAA CAC AAA CGG TTC CC (20) | 117 | DD |
| | | MS_114 | RP: CAT TCC TGG ACC CAA AAC GC (20) | | |
| 4 | *c-Met* | MS_149 | FP: GAA AGA CTT CAG CCA TCC CA (20) | 101 | DD |
| | | MS_150 | RP: GAA AGA CTT CAG CCA TCC CA (23) | | |
| 5 | *Hgf* | MS_675 | FP: GTG TGC CAA CAG GTG TAT CAG (21) | 132 | DD |
| | | MS_676 | RP: CCA AAC CCT TTT TTC ACT CCA (21) | | |
| 6 | *Pax3* | MS_111 | FP: ATT GCT CAA GGA CGC TGT (18) | 178 | DD |
| | | MS_112 | RP: TCG CTC ACT CAG GAT GCC A (19) | | |
| 7 | *Cdh15* | MS_103 | FP: ATC TAC AGC ATC CAG GGT CC (20) | 157 | DD |
| | | MS_104 | RP: TAG AGC CAC CCA AGT CCA AG (20) | | |
| 8 | *Msc* | MS_109 | FP: GAG GAC CGC TAC GAG GAC (18) | 175 | DD |
| | | MS_110 | RP: CCC AGT CCT TTG CGT TTA CC (20) | | |
| 9 | *Slit1* | MS_129 | FP: CTG CTC CCC GGA TAT GAA CC (20) | 81 | Chen Dun Neural Regen Res. 2020 |
| | | MS_130 | RP: TAG CAT GCA CTC ACA CCT GG (20) | | |
| 10 | *Slit2* | MS_131 | FP: AAC TTG TAC TGC GAC TGC CA (20) | 155 | |
| | | MS_132 | RP: TCC TCA TCA CTG CAG ACA AAC T (22) | | |
| 11 | *Slit3* | MS_133 | FP: AGT TGT CTG CCT TCC GAC AG (20) | 188 | |
| | | MS_134 | RP: TTT CCA TGG AGG GTC AGC AC (20) | | |
| 12 | *Robo1* | MS_135 | FP: GCT GGC GAC ATG GGA TCA TA (20) | 94 | |
| | | MS_136 | RP: AAT GTG GCG GCT CTT GAA CT (20) | | |
| 13 | *Robo2* | MS_137 | FP: CGA GCT CCT CCA CAG TTT GT (20) | 135 | |
| | | MS_138 | RP: GTA GGT TCT GGC TGC CTT CT (20) | | |
| 14 | *Robo3* | MS_139 | FP: CCT GTT CAA ACC CAG GAC AGC (21) | 177 | Carr Dun Plos One 2017 |
| | | MS_140 | RP: ACA CAC GGA ATC CTT GCA CC (20) | | |
| 15 | *Robo4* | MS_141 | FP: AAG CCC AGG TCC AAA CTC TG (20) | 87 | |
| | | MS_142 | RP: GTT GCG GTG AAG TTG TGG TC (20) | | |

**Supplementary Table 3. List of antibodies used**

| Antibody | Type | Manufacturer | Product No. (Clone) |
|---|---|---|---|
| Met | Monoclonal, Mouse IgG1κ | Santacruz | sc-8057 (B-2) |
| Met | Polyclonal, Goat IgG | R&D Biosystems | AF527 |
| p-Met (Tyr1234/1235) | Monoclonal, Rabbit | Cell Signaling Technology | 3077 (D-26) |
| Myosin 4 | Monoclonal, Mouse IgG2bκ | Thermo Fisher Scientific | 14-6503-82 (MF-20) |
| Slit1 | Polyclonal, Rabbit IgG | Thermo Fisher Scientific | PA5-119606 |
| Cleaved Caspase 3 (CC-3) | Polyclonal, Rabbit | Cell Signaling Technology | 9661 |
| M-Cadherin | Monoclonal, Rabbit IgG | Cell Signaling Technology | 40491 (D4B9L) |
| Musculin | Polyclonal, Rabbit IgG | Thermo Fisher Scientific | PA5-101067 |
| P-p44/42 | Monoclonal, Rabbit IgG | Cell Signaling Technology | 4370 (D13.14.4E) |
| T-p44/42 | Monoclonal, Rabbit IgG | Cell Signaling Technology | 4695 (137F5) |
| p-Akt | Monoclonal, Rabbit IgG | Cell Signaling Technology | 4060 (D9E) |
| T-Akt | Monoclonal, Rabbit IgG | Cell Signaling Technology | 4691 (C67E7) |
| Pax3 | Polyclonal, Rabbit IgG | Thermo Fisher Scientific | PA1107 |
| Neurofilament -L (C28E10) | Monoclonal Rabbit IgG | Cell Signaling Technology | 2837 (C28E10) |
| a-Myosin (Skeletal, Fast) − Alkaline Phosphatase conjugated | Monoclonal Mouse IgG1 | Sigma | A4335 (MY-32) |
| Laminin | Polyclonal Rabbit | Sigma | L9393 (LAMA1) |
| β-Actin | Monoclonal Mouse IgG2b | Cell Signaling Technology | 3700 (8H10D10) |
| GAPDH | Monoclonal Mouse IgG1 | Thermo Fisher Scientific | MA5-15738 |
| α–Tubulin | Monoclonal, Mouse IgG1κ | Thermo Fisher Scientific | 62204 (DM1A) |
| Cy3 conjugated Donkey α-Goat | Donkey IgG (H+L) | Jackson ImmunoResearch Laboratories | 705-165-147 |
| Cy3 conjugated Goat α-Rabbit | Goat IgG (H+L) | Jackson ImmunoResearch Laboratories | 111-165-144 |
| Cy2 conjugated Donkey α-Rabbit | Donkey IgG (H+L) | Jackson ImmunoResearch Laboratories | 711-225-152 |
| Cy2 conjugated Goat α-Mouse | Goat IgG (H+L) | Jackson ImmunoResearch Laboratories | 115-225-146 |
| HRP conjugated Goat α-Mouse | Goat IgG (H+L) | Jackson ImmunoResearch Laboratories | 115-035-003 |
| HRP conjugated Goat α-Rabbit | Goat IgG (H+L) | Jackson ImmunoResearch Laboratories | 111-035-144 |

**Supplementary Figure 2**

Dispersion plot

Dispersion plots help estimate and visualize variability in gene expression across the replicate samples of the compared groups (cMet$^{KO}$ and WT embryos). Since dispersion is a measure of variability in gene expression relative to the mean, dispersion for each gene is estimated using the read counts of the gene in all replicates. These initial gene-wise dispersion estimates/variances are then plotted as a function of their respective mean expression levels resulting in a dispersion plot, where each gene is indicated by a black dot. In the dispersion plot, a curve (shown as a red line) is fitted to the gene-wise dispersion estimates representing the expected dispersion value for genes. The dispersion estimates are then adjusted by shrinking the initial gene-wise dispersion estimates, using appropriate statistical methods, towards the fitted curve (red line). The final adjusted dispersion values so obtained are represented by blue dots in the dispersion plot. The genes represented as black dots surrounded by blue circles in the dispersion plot, are genes that that have extremely high dispersion, possibly because of their higher variability or technical issues, and shrinking their dispersion estimates towards the curve (red line) could result in false positives. Thus, dispersion shrinkage helps reduce false positives (incorrectly identified DEGs) in the differential expression analysis. The dispersion plots for FLB and HLB datasets for the compared groups in this study are typical as they show a downward trend where genes with high expression (i.e. high mean counts) have lower dispersion values, and the genes with low expression (i.e. low mean counts) have higher dispersion estimates.

PCA plot

Principal component analysis (PCA) helps analyse high dimensional gene expression data from RNAseq datasets. A PCA plot reduces or transforms this complex dimensionality to a small number of principal components (PCs or dimensions), enabling a 2D visualization of relationship between biological samples based on their gene expression patterns. Samples having similar gene expression patterns cluster together on the PCA plot, suggesting their biological and functional similarity. The first and the second principal components (PC1 and PC2) are axes in the PCA plot that explain specific percentage of variance of the total variation in the gene expression data. Typically, PC1 captures larger percentage of the total variance compared to PC2.
